# Sex differences in nucleus accumbens core circuitry engaged by binge-like ethanol drinking

**DOI:** 10.1101/2024.08.15.608144

**Authors:** Amy E. Chan, Justin Q. Anderson, Kolter B. Grigsby, Bryan E. Jensen, Andrey E. Ryabinin, Angela R. Ozburn

## Abstract

Growing parity in Alcohol Use Disorder (AUD) diagnoses in men and women necessitates consideration of sex as a biological variable. In humans and rodents, the nucleus accumbens core (NAcc) regulates alcohol binge drinking, a risk factor for developing AUD. We labeled NAcc inputs with a viral retrograde tracer and quantified whole-brain c-Fos to determine the regions and NAcc inputs differentially engaged in male and female mice during binge-like ethanol drinking. We found that binge-like ethanol drinking females had 129 brain areas with greater c-Fos than males. Moreover, ethanol engaged more NAcc inputs in binge-like ethanol drinking females (as compared with males), including GABAergic and glutamatergic inputs. Relative to water controls, ethanol increased network modularity and decreased connectivity in both sexes and did so more dramatically in males. These results demonstrate that early-stage binge-like ethanol drinking engages brain regions and NAcc-inputs and alters network dynamics in a sex-specific manner.

**Graphical Summary:**
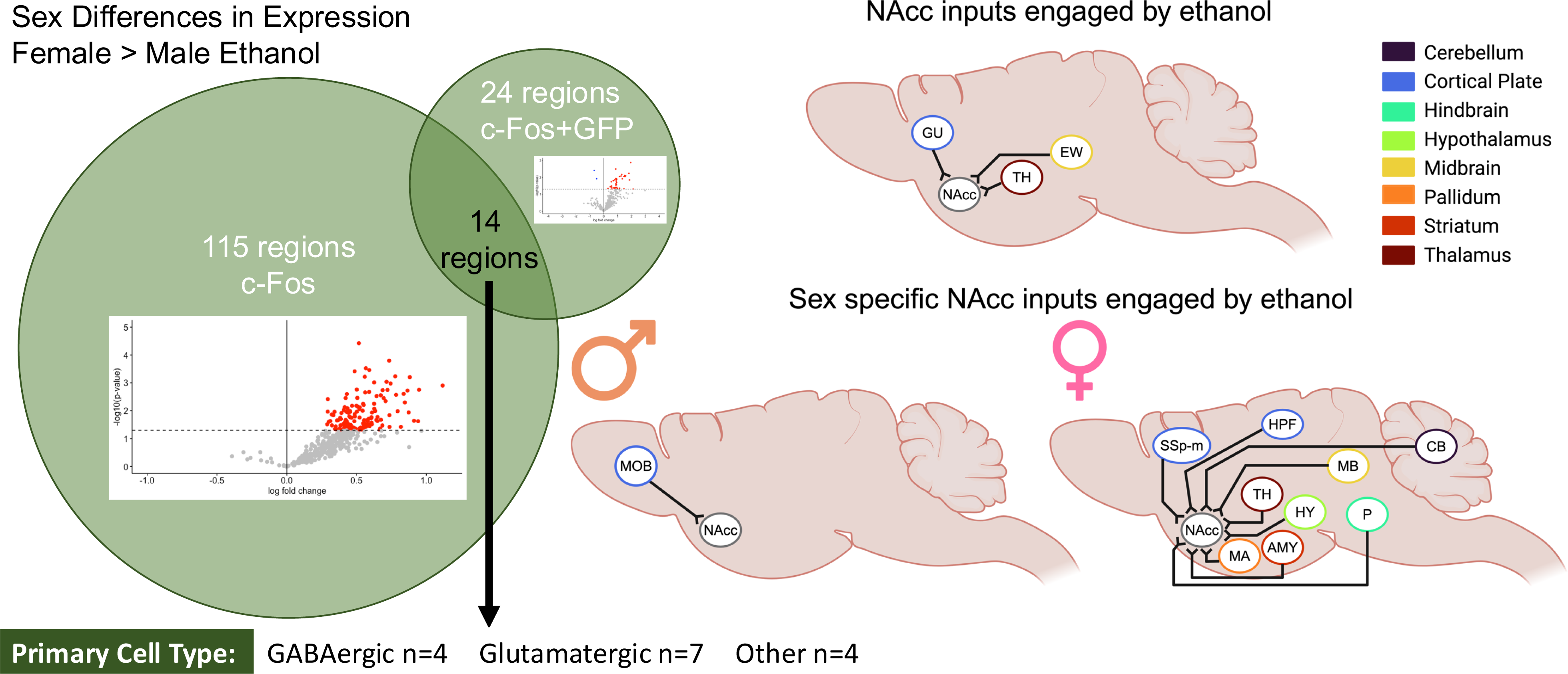
Binge-like ethanol drinking engages more regions and NAcc inputs in female relative to male mice. *Left:* Comparison of regions with both greater c-Fos expression and c-Fos+GFP colocalization in female relative to male ethanol drinking mice. *Right:* NAcc inputs engaged by binge-like ethanol drinking compared to water drinking mice, sex collapsed. Thalamic (TH) regions include left parasubthalamic nucleus, left anteromedial nucleus of the thalamus, left central medial nucleus of the thalamus, left medial group of the dorsal thalamus, left subparafascicular nucleus, left peripeduncular nucleus, and right paraventricular nucleus of the thalamus. EW, Edinger-Westphal nucleus; GU, gustatory areas. *Bottom Middle:* NAcc inputs with greater engagement in male than female ethanol drinking mice (left and right main olfactory bulbs (MOB). *Bottom Right:* NAcc inputs with greater engagement in female than male ethanol drinking mice. Amygdala (AMY) regions include left anterior amygdalar area and left intercalated amygdalar area. Hippocampal (HPF) regions include right dentate gyrus, right Field CA1, right Field CA2, and right Field CA3. Hypothalamic (HY) regions include left and right lateral hypothalamic area, left and right periventricular hypothalamic nucleus, preoptic part, right dorsomedial nucleus of the hypothalamus, right posterior hypothalamic nucleus, and right ventromedial hypothalamic nucleus. Midbrain (MB) regions include left and right midbrain reticular nucleus, retrorubral area, left and right superior colliculus, motor related, left nucleus of the brachium of the inferior colliculus, left nucleus of the posterior commissure, left olivary pretectal nucleus, left posterior pretectal nucleus, left superior colliculus, sensory related, and right substantia nigra, reticular parts. Pontine (P) regions include left superior central nucleus raphe, left supratrigeminal nucleus, right nucleus raphe pontis, right pontine reticular nucleus, and right superior olivary complex. Thalamic (TH) regions include left and right lateral dorsal nucleus of the thalamus, right dorsal part of the lateral geniculate complex, right lateral posterior nucleus of the thalamus, and right parasubthalamic nucleus. CB, Cerebellum; MA, magnocellular nucleus; SSp-m, primary somatosensory cortex, mouth; Created with BioRender.com. See **Table S18** for additional information.

## Introduction

Binge alcohol drinking, defined as drinking to intoxicating blood alcohol concentrations (>80mg/dL), is a risk factor for developing Alcohol Use Disorder (AUD; [1]). While alcohol use and AUD diagnoses are more prevalent in men than women, over the past 20 years this gender gap has drastically narrowed [2, 3]. Women have more severe health outcomes in response to alcohol, including increased risk of developing breast-cancer and greater death rates [4–7]. Moreover, current treatments for AUD are not always equally effective in both sexes [8, 9]. Considering the growing evidence of sex differences in the neurobiology of AUD, it is imperative to consider sex as a biological variable (SABV, [10–13]).

Much of our understanding of the neurocircuitry underlying addictive disorders comes from studies with male subjects. The nucleus accumbens core (NAcc) is a region that is critically involved in all stages of AUD [14]. In treatment resistant male patients with AUD, nucleus accumbens (NAc) deep-brain stimulation (DBS) reduced alcohol craving and promoted lower intake or abstinence [15, 16]. NAcc DBS also reduced ethanol intake in male rats [17]. This evidence supports that the NAcc is a clinically relevant target for the treatment of AUD and that its engagement in AUD can be modeled in rodent studies.

The rodent NAcc exhibits sex differences in response to ethanol, and behavioral outcomes following manipulations of NAcc activity. Male and female C57BL/6J (B6) mice exhibit distinct transcriptional responses in the NAc following limited access binge-like ethanol drinking [18]. Chemogenetic NAcc inhibition increased binge-like ethanol intake in female B6 mice, but decreased intake in male mice [19–21]. Conversely, NAcc stimulation decreased binge-like ethanol intake in females, and increased intake in males [19–21]. This evidence underscores the importance of explicitly addressing SABV while identifying neural mechanisms underlying risk for AUD.

Studies using c-Fos, an immediate early gene used as a proxy for cellular activity, have informed our understanding the brain regions engaged by alcohol and drugs of abuse. Burnham & Thiele (2017) identified several regions with elevated c-Fos after binge-like ethanol intake in male B6 mice, and other studies have identified regions with anatomical projections to the NAcc that are engaged by ethanol [22–30]. However, it is currently unknown if these projections are engaged during binge-like drinking.

This study sought to fill critical gaps in the literature by determining whether early-stage binge-like ethanol drinking engages brain regions, networks, and NAcc inputs in a sex-specific manner. We employed whole-brain quantification of c-Fos to identify novel regions and circuits of interest, and understand changes in network scale dynamics [31]. Overall, we found that female mice had both more regions and NAcc inputs engaged than males following binge-like ethanol intake, and, relative to same sex water drinking controls, ethanol reduced network connectivity more greatly in males.

## Results

### Male and female mice drink ethanol to intoxication

Male and female mice went through a 4-day Drinking-in-the-Dark (DID) procedure with 20% ethanol or water as previously described (**Figure 1A**, [32]). There was a significant effect of sex on 2hr ethanol intake, where females displayed higher intake than males (**Figure 1B**). Most mice drank to intoxication and there was no effect of sex on 4hr ethanol intake or the achieved blood ethanol concentration (BEC, **Figure 1C-D**). There was no effect of sex on water intake (**Figures 1E-F**).

**Figure 1.**
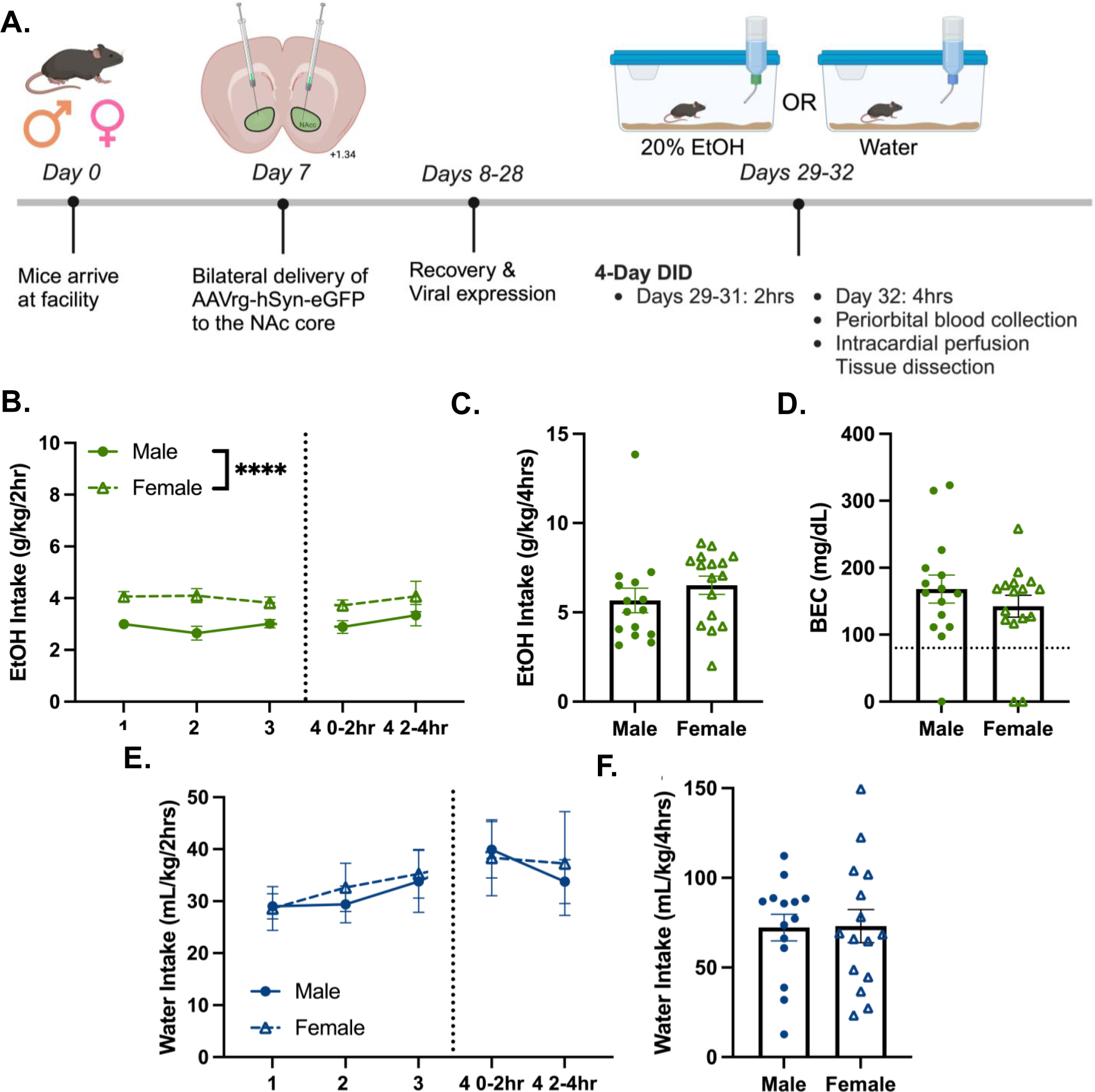
Male and female mice drink ethanol to intoxication. (A) Experimental timeline. Created with BioRender.com (B) 2hr ethanol intake over 4-day DID (n=15-16/sex; day 1-3 and day 4 0-2hr intake, F_(1,29)_ = 25.20; ****p<0.0001) (C) Day 4, 0-4hr ethanol intake. (D) BEC after 4hr ethanol intake, mean BECs: 155mg/dL ±0.13 SEM. (E) 2hr water intake over 4-day DID (n=15/sex). (F) Day 4 0-4hr water intake. Ethanol denoted by green symbols; water denoted by blue symbols. Closed circles denote males; open triangles denote females. Data shown as mean ± SEM.

### Sex and fluid drive differences in c-Fos expression

Hierarchical clustering of the average c-Fos expression per region in each sex and fluid group revealed that while both water drinking groups cluster together, the ethanol groups are on separate branches of the dendrogram (**Figure 2A**), indicating dissimilarity in c-Fos expression. Principal Components Analysis (PCA) revealed that fluid (ethanol or water) was significantly associated with ∼7.7% of explained variance in the data through associations with Principal Components (PCs) 5 (∼3.2%), PC6, (∼2.6%), and PC9 (∼1.9%; **Figure 2B**). While sex was not significantly associated with any of the top 10 PCs, there was a trending association with PC1 (∼41.8%) and PC7 (∼2.5%). Together, this accounts for 44.3% of the total explained variance in the data, more than any other factor considered. A sex by fluid interaction was not significantly associated with any of the top 10 PCs but had a trending association with PC9 (1.94%; **Table S2**).

**Figure 2.**
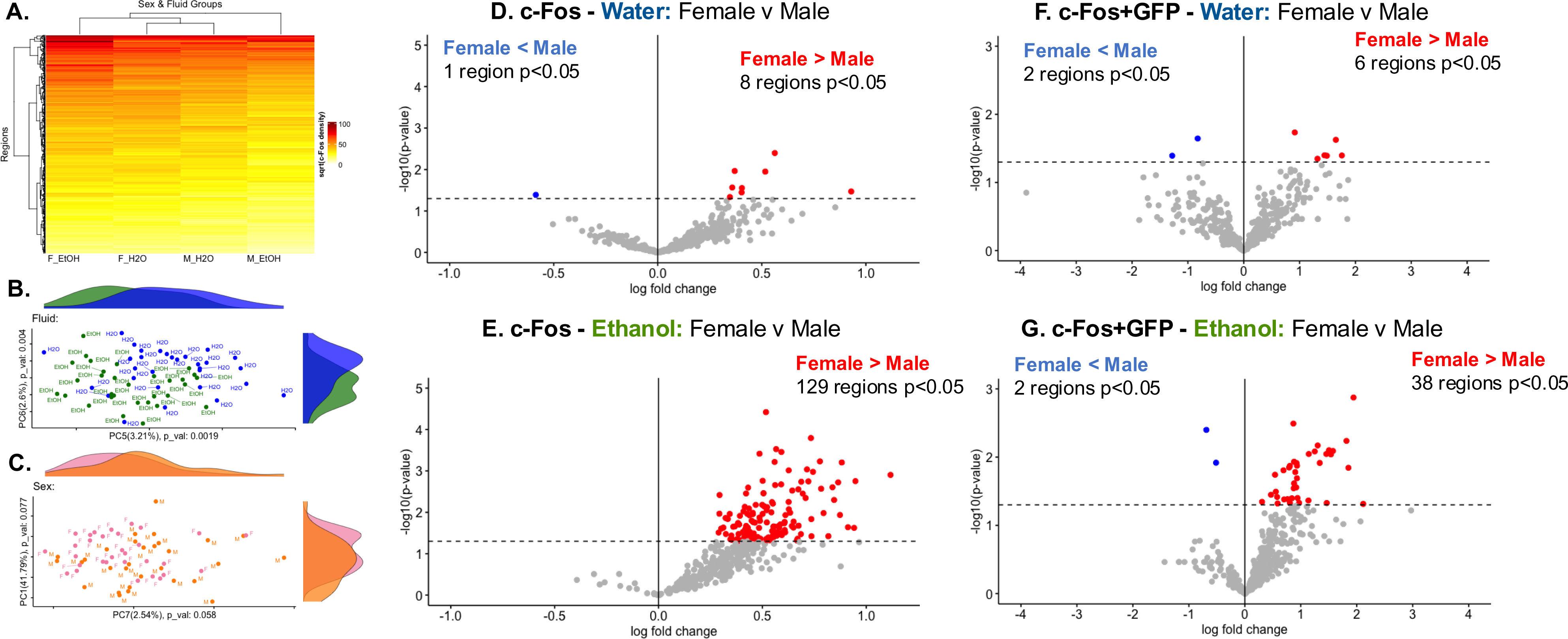
Sex and fluid drive differences in c-Fos expression and NAcc input engagement. (A) Hierarchically clustered heatmap showing average c-Fos expression [reduced, square root transformed density (cells/mm^3^)]. Y-axis clusters regions by similarity, x-axis clusters sex and fluid subgroups by similarity. (B & C) Top 2 PCs and proportion of explained variance associated with fluid (PC5 - 3.2%, PC6 - 2.6%, PC9 - 1.9%; 1-way ANOVA, p<0.05), or sex (PC1 - 41.8%, PC7 - 2.5%). See **Table S2** for further information. (D-G) Volcano plots comparing c-Fos expression (D&E, n=15-16/sex/fluid) or c-Fos+GFP colocalization (F&G, mice with bilateral NAcc GFP viral expression, n=7-14/sex/fluid) between sexes of the same fluid group. Dotted line at –log10(p-value)=1.3 denotes threshold for significance (t-test, p<0.05). Individual regions are plotted, color corresponds to the direction of log-fold change and significance. (D) One region had significantly less c-Fos expression in female than water drinking mice. 8 regions had significantly greater c-Fos expression in female than male water drinking mice. (E) No regions had significantly less c-Fos expression in female than male ethanol drinking mice. 129 regions had significantly greater c-Fos expression in female than male ethanol drinking mice. (F) 2 regions had significantly less c-Fos+GFP colocalization in female than male water drinking mice. 6 regions had significantly higher c-Fos+GFP colocalization in female than male water drinking mice. (G) 2 regions had significantly less c-Fos+GFP colocalization in female than male ethanol drinking mice. 38 regions had significantly greater c-Fos+GFP colocalization in female than male ethanol drinking mice. See **Tables S3 and S5** for further information.

### Females have greater regional engagement than males during binge-like ethanol drinking

Given that both sex and fluid drive variance in the data, we determined which regions had differential c-Fos expression in a sex-collapsed and sex specific manner. In sex-collapsed data, 6 regions had significantly greater c-Fos expression in ethanol compared to water drinking mice. This included the left and right Edinger-Westphal nucleus (EW; **Figure S3A**), a region highly sensitive to experimentally delivered and voluntarily consumed ethanol [33, 34]. Next, we determined whether there were differences in c-Fos expression between male and female mice within a fluid group. 8 regions had greater c-Fos expression in female than male mice water drinking mice, including the right ventral premammillary nucleus and left medial amygdala regions (**Figure 2D**) Both regions have sexually dimorphic functions and cellular composition (for review see [35, 36]). 1 region had less c-Fos expression in female than male mice water drinking mice, the left dorsal tegmental nucleus.

These results demonstrate subtle baseline differences in c-Fos density between males and females. 129 regions had significantly greater c-Fos expression in female than male ethanol drinking mice (**Figure 2E**). Notable regions include the right nucleus accumbens, bed nucleus of the stria terminalis, pallidum (caudal region), right infralimbic area, the left agranular insular area (dorsal and ventral parts), and the left and right claustrum. No regions had less c-Fos expression in female than male ethanol drinking mice. Together this indicated that the female mouse brain was engaged to a greater extent following binge-like ethanol drinking relative to the male mouse brain (**Figure S3** and **Table S3** for additional comparisons).

### Females have great engagement of NAcc inputs than males during binge-like ethanol drinking

Similarly to the c-Fos data, we determined which NAcc inputs (through quantification of c-Fos+GFP colocalized cells) had differential engagement in a sex-collapsed and sex specific manner. In sex-collapsed data, 10 regions had significantly greater c-Fos+GFP colocalization in ethanol compared to water drinking mice, including the left EW (**Figure S7A**), a known source of projections to the NAcc [37]. Next, we determined whether there were differences in c-Fos+GFP colocalization between male and female mice within a fluid group. 6 regions had greater c-Fos+GFP colocalization in female than male water drinking mice, including parts of the pretectal nucleus in both hemispheres. 2 regions had less c-Fos+GFP colocalization in female than male water drinking mice (both were left hemisphere hypothalamic nuclei; **Figure 2F**). These results elucidate subtle baseline differences in NAcc input engagement between male and female water drinking mice. 38 regions had significantly greater c-Fos+GFP colocalization in female than male ethanol drinking mice (**Figure 2G**). This included 14 regions that also had greater c-Fos density in ethanol drinking female than male mice, notably, the left anterior amygdalar area, right hemisphere hippocampal regions (dentate gyrus, Field CA1 and CA3) and nucleus raphe pontis, and 6 hypothalamic nuclei across both hemispheres. 2 regions had less c-Fos+GFP colocalization in female than male ethanol drinking mice, the left and right main olfactory bulbs. Together, this demonstrates that more NAcc inputs in females are engaged relative to males following binge-like ethanol drinking (**Figure S7** and **Table S5** for additional comparisons).

### Network analyses of c-Fos density

#### Method 1: Correlation matrices

##### Sex and fluid drive differences in network structure

To determine the impact of sex and fluid on network structure and dynamics, correlation matrices were constructed from c-Fos expression for each sex and fluid group, similarly to prior analyses [38–41]. The female water network had a greater number of modules than the male water network, but female and male ethanol networks had the same number of modules (male water n=5; female water n=9; male ethanol n=12; female ethanol n=12; **Figure 3A-D**). Ethanol networks had more modules than sex-matched water networks, at all tree-cut heights of 90% and lower, suggesting this is a robust feature of the data (**Figure 3I**). The male water network had the least modularity of any group. Together, this indicated 1) differences in baseline network structure between water drinking male and female mice, and 2) that network structure is further perturbed by binge-like ethanol intake.

**Figure 3:**
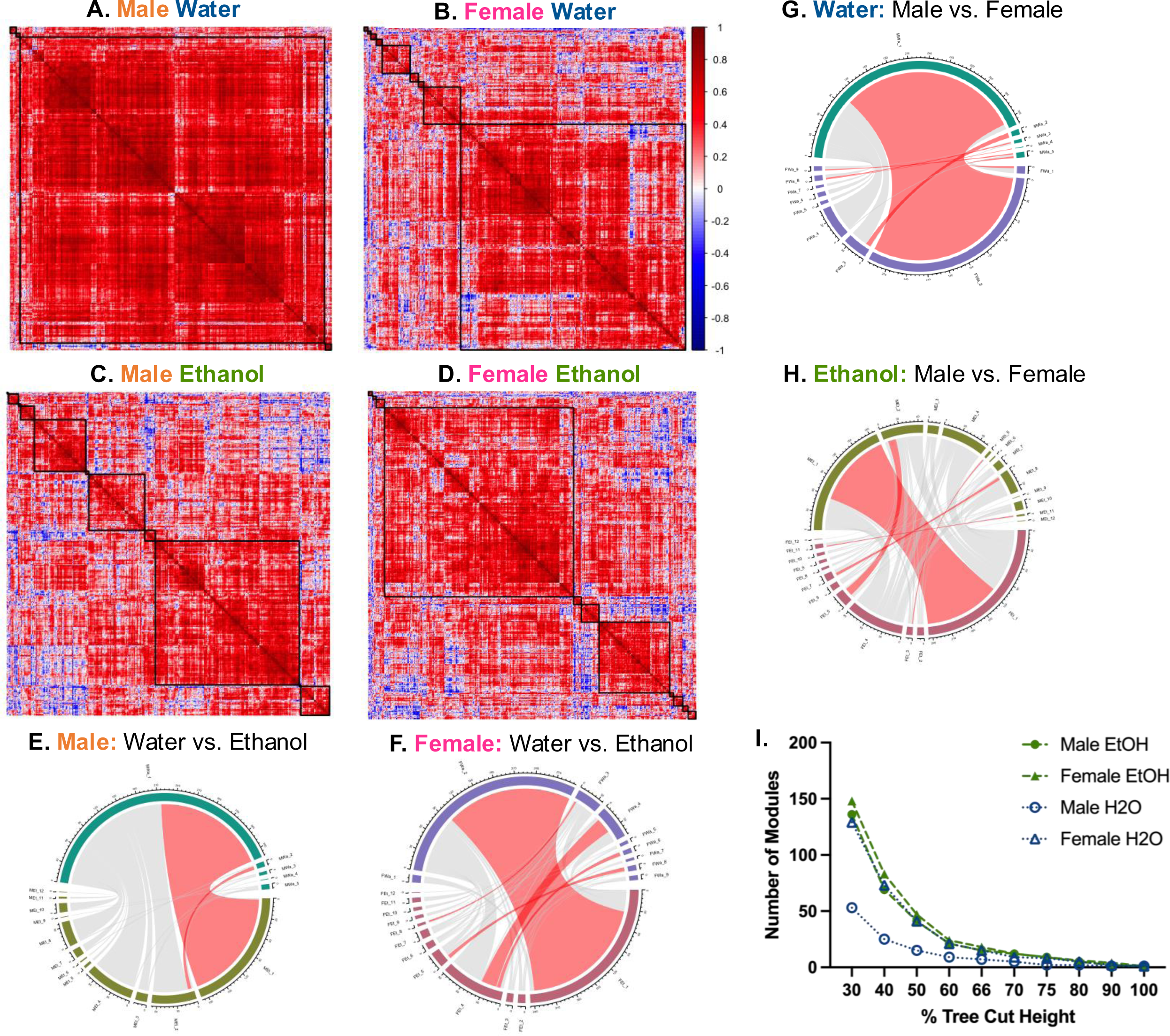
Sex and fluid determine network module number, size, and composition. (A-D) Hierarchically clustered heatmaps of Pearson’s correlation matrices [R, square root transformed c-Fos density (cells/mm^3^)] for each sex and fluid group. Black squares denote the modules determined when the cluster dendrogram was cut at 70% tree height. (A) Male water (n=5 modules). (B) Female water (n=9 modules). (C) Male ethanol (n=12 modules). (D) Female ethanol (n=12 modules). (E-H) Chord diagrams comparing number of regions in each module within sex (E&F) or fluid (G&H) groups. Red chords indicate modules with a significant number of regions that overlap between networks (fisher’s test, fdr<0.05). (E) Significant overlap of regions between MWa1 and MEt, and MWa2 and MEt2. (F) Significant overlap of regions between FWa2 and FEt1, FWa3 and FEt8, FWa4 and FEt4, FWa6 and FEt4, and FWa8 and FEt5. (G) Significant overlap of regions between MEt1 and FEt1, MEt2 and FEt5, MEt5 and FEt3, MEt8 and FEt7, and MEt11 and FEt11. (H) Significant overlap of regions between MWa1 and FWa2, MWa2 and FWa3, MWa3 and FWa8, MWa4 and FWa8, MWa5 and FWa1, and MWa5 and FWa9. (I) Number of modules by percent tree cut height. Ethanol denoted by green, filled symbols, water denoted by open, blue symbols. Females denoted as triangles, males denoted by circles. MWa, male water; MEt, male ethanol; FWa, female water; FEt, female ethanol. See **Tables S6, S8, S10,** and **S12** for further information.

##### Sex and fluid drive differences in network composition

Next, we determined whether any modules had overrepresentation of regions with differential c-Fos expression (greater c-Fos in female relative to male ethanol drinking mice). Only the male ethanol module 1 had significant overrepresentation of regions with differential c-Fos expression [72 regions overlap, **Table S10**]. This indicates that regions with differential c-Fos cluster within a single male ethanol module but are dispersed among several modules in all other networks (**Figures S8-S9, and Tables S6-S13** for additional comparisons).

###### Fewer modules are preserved between male networks than female networks

To compare module composition between networks, we determined whether there was module preservation between sex-matched or fluid networks. Only 2 modules significantly overlapped between male networks (**Figure 3E**), whereas 5 modules significantly overlapped between female water and ethanol networks (**Figure 3F**). Together, this indicates that ethanol increased modularity and network dissimilarity to a greater extent in males. 6 modules significantly overlapped between the male and female water drinking networks (**Figure 3G**). Similarly, 5 modules significantly overlap between the male and female ethanol networks (**Figure 3H**), indicating aspects of network structure are preserved between males and females of the same fluid group.

##### Sex and fluid drive differences in network connectivity

###### Ethanol reduced intramodular connectivity in males

Further, we explored whether the intramodular correlations between regions in each network’s largest module were influenced by sex and fluid (**Figure S9I**). The largest male water module had significantly greater correlation coefficients than modules of all other groups. The largest male ethanol module had significantly greater correlation coefficients than the largest modules in the female ethanol and water networks. Average intramodular correlation values were not significantly different between the female ethanol and water modules. This demonstrates that ethanol reduces intramodular connectivity in males but not females, and that the male modules had greater intramodular connectivity than female modules. Further analysis indicated that regions from the cortical plate was significantly overrepresented in the largest modules of each network (**Tables S6, S8, S10, S12**). Therefore, differences in intramodular correlation strength between sex and fluid networks may also reflect differences connectivity of regions within the cortical plate.

###### Ethanol reduced the connectivity of more regions in males than females

We determined whether regional connectivity changed between water and ethanol networks of the same sex, and if the direction of change or regions changed were similar for both sexes. 100 regions had less connectivity in the male ethanol relative to the water network (**Table S14**). Of these differentially wired regions, 42 were also regions with differential c-Fos expression. 33 regions were differentially wired between female water and ethanol networks (**Table S15**). 12 regions had greater connectivity in the female ethanol relative to the female water network, including 3 regions with differential c-Fos expression. 21 regions had less connectivity in ethanol relative to water drinking females, including 11 regions with differential c-Fos expression.

In male and female networks, 5 regions had reduced connectivity in response to ethanol, including the right thalamus (anteromedial, anteroventral, interanterodorsal, and rhomboid nuclei) and right caudoputamen, suggesting that ethanol reduced connectivity of these regions similarly. Only 1 region had increased connectivity in the female ethanol network, but reduced connectivity in the male ethanol network, the left ventral posterior complex of the thalamus (VP). In the female ethanol network, connections between the left VP and 8 other regions were increased compared to the female water network, including parts of the visual system (left anteromedial, posteromedial and primary visual areas, right anterolateral and lateral visual areas) and the left dorsal and lateral agranular areas of the retrosplenial cortex. In the male ethanol network, the left VP had reduced connectivity to 16 regions compared to the male water network, including the right hemisphere anterior amygdala, Field CA3, substantia nigra, *pars reticulata* and *pars compacta*, main olfactory bulb, and ventral tegmental nucleus. This evidence shows that ethanol reduced connectivity more in male than female networks, changed the connectivity of different regions in male and female networks, and altered connectivity of the same region in different directions.

###### Ethanol increased inter-modular connectivity to a greater extent in males

Next, we determined whether ethanol changed the way regions (nodes) were connected both within and between modules, similarly to prior work [38, 39, 42]. In male and female networks, ethanol changed the number of each type of node (**Figure S8D**). In males, ethanol dramatically decreased in the number of ultraperipheral nodes and provincial hubs and increased all other types of nodes. Similarly in females, ethanol decreased the number of provincial hubs and increased the number of hub and non-hub connector nodes. There was a significant difference in node and hub composition between male and female ethanol networks. The male ethanol network had a greater number of hub and non-hub connector nodes, while the female ethanol network had more peripheral nodes and provincial hubs. Together this indicated that ethanol increased inter-module connectivity by increasing the proportion of connector nodes in a network, but to a greater extent in males (**Tables S7, S9, S11, S13**).

###### Male and female ethanol networks have unique hubs modules

We evaluated whether any hubs were shared between networks to determine regions that may be important drivers of ethanol drinking behaviors. Male ethanol and water networks shared more hubs (29 hubs) that shared between female ethanol and water networks (13 hubs). The hubs shared between ethanol and water networks tended to be provincial hubs in the water networks, but connector hubs in the ethanol networks. While these hubs may not be unique to ethanol drinking, ethanol did alter their function within the network. 50 regions were hubs in the male ethanol, but not male water network. 69 regions were hubs in the female ethanol, but not female water network. 16 hubs were shared between ethanol networks, 9 of which are not hubs in either water network. In both ethanol networks, the 9 non-water hubs were primarily provincial hubs in the female ethanol network, but connector hubs in the male ethanol network. Of these hubs, 7 were regions that had differential c-Fos expression, including the right ventral anterior cingulate area, right presubiculum, and left subiculum. In males, all but 1 of these regions were connector hubs, while in females 3 of these were connector hubs and 4 were provincial hubs. Thus, while these hubs are associated with ethanol intake in both sexes, they have different network connections.

Hubs have strong connections with multiple regions, through which they can influence ethanol drinking behaviors. The 27 unique hubs in the male ethanol network included 3 regions with differential c-Fos expression (left hemisphere anterior amygdalar area, cuneiform nucleus, primary somatosensory area, trunk). 40 hubs were unique to the female ethanol network. Of these hubs, 9 were also regions with differential c-Fos expression, including the insular cortex (left ventral, right posterior subdivisions), and the right periventricular hypothalamic nucleus, preoptic part (PVpo). The female ethanol network had more unique module hubs that were regions with differential c-Fos expression than the male ethanol network, suggesting that these hubs could drive ethanol drinking behaviors in a sex-specific manner (**Tables S6, S8, S10, S12**).

### Method 2: WGCNA –Sex and ethanol intake are associated with several network modules

Weighted gene co-expression network analysis (WGCNA) was employed to explore additional network characteristics (**Figure 4A, Table S18**). 8 modules were associated with sex (**Figure 4B**). No modules were associated with fluid or sex by fluid interaction. The module eigenregions (MEs) of ethanol drinking mice in 5 modules were significantly correlated with 4hr ethanol intake (day 4), and 9 modules were significantly correlated with total 4-day ethanol intake. No modules were significantly correlated with BEC.

**Figure 4.**
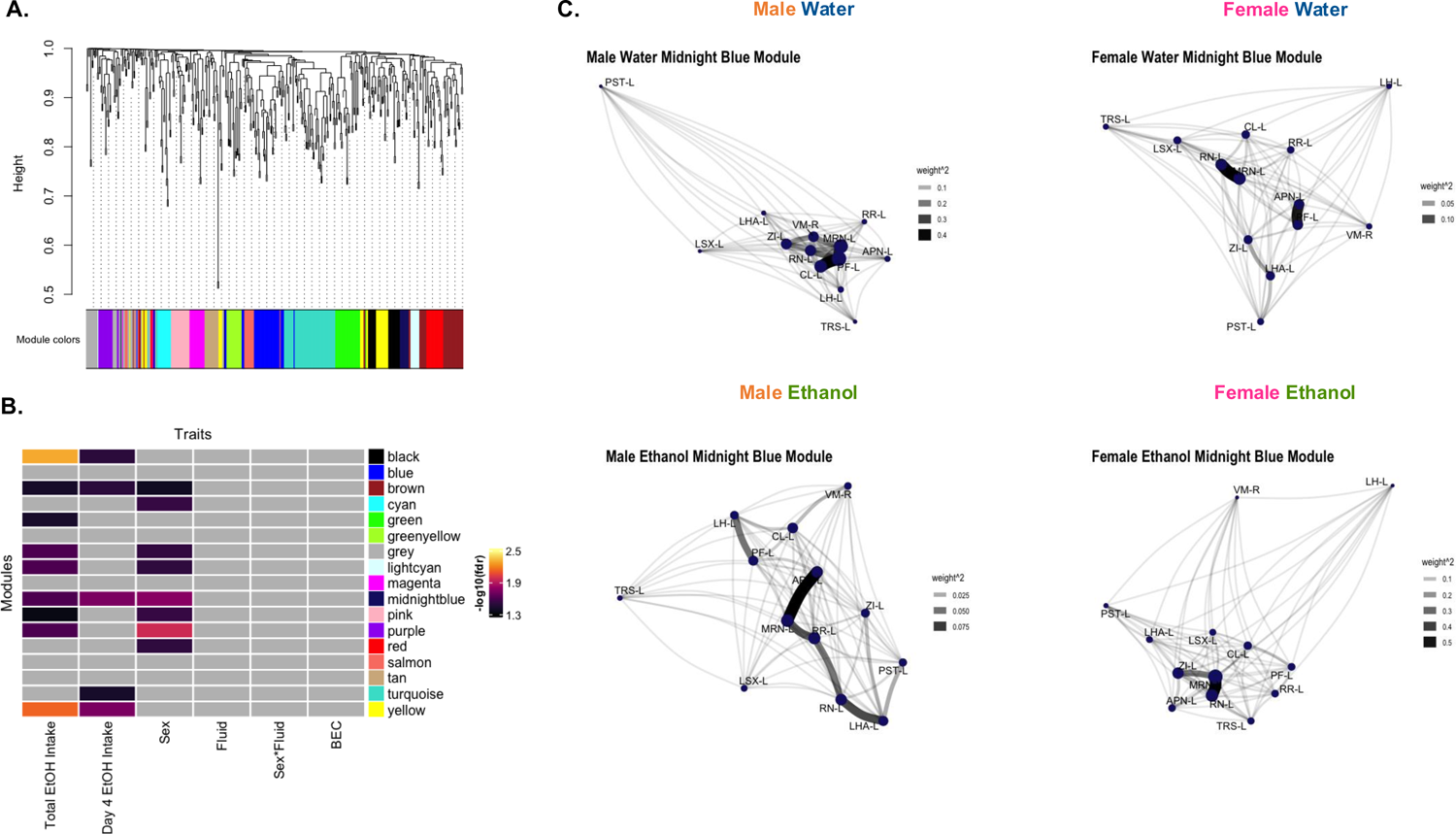
WGCNA identifies modules associated with sex and ethanol intake. (A) WGCNA created with c-Fos expression [reduced, square root transformed density (cells/mm^3^); n=14-16/sex/fluid, 1 male water mouse excluded as an outlier]. Dendrogram clustering regions by module eigenregion values (MEs) and the module color for each region. (B) Relationships between modules and traits. Total EtOH Intake (sum of ethanol intake on all 4 days) and day 4 ethanol (EtOH) intake was significantly correlated with MEs of ethanol drinking mice in several modules (Student’s asymptotic p-value, fdr <0.05). Sex was significantly associated with MEs of several modules (2-way ANOVA, fdr’s<0.05). Gray squares indicate non-significant p-values. (C) Network graphs of relationship between regions in the midnight blue module in each sex and fluid group. Graphs created from adjacency matrices for each sex and fluid group. Node size denotes importance of node in network (larger nodes are hubs). Opacity and width of links denotes strength of connections. See **Table S1** and **S16** for further information.

The 2 modules associated with sex and both metrics of ethanol intake likely influenced ethanol intake differently in males and females. The brown module was comprised of 38 regions and had significant overrepresentation of regions in the midbrain and right hemisphere (**Table S17**). 5 of the 7 hubs in this module were regions with differential c-Fos expression, including the right midbrain reticular nucleus and right periaqueductal gray. The midnight blue module contained 13 regions, had significant overrepresentation of regions in the left hemisphere, and both module hubs were regions with differential c-Fos expression (left midbrain reticular nucleus and red nucleus). Network graphs of regions in this module (**Figure 4C)** illustrate that the hubs, type, and strength of connections between these regions varies by sex and fluid group.

The composition of nodes and hubs in the WGCNA network was significantly different than all the correlation matrix networks, likely because data from all sex and fluid groups were included in the WGCNA network, whereas correlation matrix networks were created for each sex and fluid group separately. Similarly to the ethanol correlation networks, most non-hub regions were provincial nodes, and most hubs were connector hubs. The WGCNA network had fewer non-hub and hub kinless nodes than the correlation matrix networks.

## Discussion

We used whole-brain c-Fos quantification and a fluorescent retrograde tracer to identify the regions and NAcc specific inputs engaged by early-stage binge-like ethanol drinking. Further, we leveraged the depth of the whole brain c-Fos data to identify network characteristics altered by binge-like drinking. The present work demonstrated that early-stage binge-like ethanol drinking engages more regions and NAcc inputs in female relative to male mice. We also found that ethanol increases modularity and decreases intramodular connectivity to a greater extent in males. The number of significant sex differences in these findings cannot be attributed to ethanol intake or BECs, as male and female mice consumed similar amounts of ethanol and had similar BECs at the time of brain collection.

The most striking findings of this study were how differently ethanol impacted the regions engaged and the network characteristics of females compared to males. No regions had significantly greater engagement in males relative to females, but 129 regions had greater engagement in female relative to male ethanol drinkers. These regions were from different anatomical categories with the greatest representation of the cortical plate (40 regions), thalamus (20 regions), and hypothalamus (20 regions). Therefore, ethanol increased regional engagement across the brain more in females than males. This finding was echoed by the differential wiring analysis which identified the left ventral posterior complex of the thalamus (VP) as a candidate for future investigation. The left VP was the only region that had increased connectivity in female, but reduced connectivity in male ethanol drinkers, relative to same-sex water drinkers. Indicative of this, the left VP was a provincial hub in the female ethanol network, but a non-hub connector node in the male ethanol network. This region processes somatosensory information, is important for shifting between behavioral states, and is currently a target for DBS to treat neuropathic pain [43–45].

Female ethanol drinking mice also had greater engagement of NAcc inputs. These 38 NAcc inputs were from all different parts of the brain, with the strongest representation from the midbrain (10 regions) and hypothalamus (9 regions). 14 of these NAcc inputs were also regions with greater c-Fos expression in female relative to male ethanol drinking mice, including 7 excitatory/glutamatergic regions, 4 inhibitory/GABAergic regions, a serotonergic region, a dopaminergic region, and 1 region with mixed cell types (**Graphical Summary, Table S18**). This finding indicated that during binge-like ethanol drinking, the female NAcc integrates information from a wide variety of inputs. In male ethanol drinking mice, only the olfactory bulbs, which send excitatory inputs to the NAcc [46, 47], were more engaged by ethanol relative to females. This observation could indicate that ethanol’s olfactory signature is processed differently in males and females. The large difference in number and types of NAcc inputs engaged by binge-like drinking could explain why chemogenetic manipulations of the NAcc impacted drinking behavior in a sex specific manner [19–21]. The present results also suggest different circuit mechanisms control binge-like drinking in males and females.

Another region of potential interest for future studies is the periventricular hypothalamic nucleus, preoptic part (PVpo). The right PVpo was a hub in the female ethanol network and the left PVpo was a hub in a WGCNA module related to sex and ethanol intake. Both the right and left PVpo were regions with differential c-Fos expression and NAcc inputs with greater engagement in female than male ethanol drinking mice. This region releases somatostatin to the median eminence and is involved in body fluid homeostasis [48, 49]. One study found a transient increase in cocaine- and amphetamine-regulated transcript peptide (CART) immunoreactivity in the PVpo 24 hours into ethanol withdrawal in male Sprague-Dawley rats [50]. Another study in male rats found no effect of ethanol withdrawal on β-endorphin expression in the PVpo [51]. Future work can address whether the PVpo-to-NAcc circuit alters drinking in sex-specific ways, and identify the signaling molecules released (i.e. glutamate, GABA, neuropeptides etc., similarly to [52]).

### Ethanol increases modularity in females, males are responsive to number of drinking days

Thus far, five studies have used whole-brain c-Fos expression to evaluate brain networks engaged in models for dependence and withdrawal or escalation of ethanol intake. An understanding of brain network structure provides information about the system, rather than activity of regions in isolation [53, 54]. In male mice, ethanol dependence reduced modularity, and there were modest reductions in modularity in non-dependent, ethanol exposed male mice compared to naïve controls [39]. Another study in dependent male mice demonstrated that individual variability in ethanol intake and withdrawal state also alters network structure [55]. In agreement, male ethanol drinking mice had lower modularity compared to water drinking males after an every other day intermittent ethanol two-bottle choice drinking [41]. However, in this same study, ethanol drinking female mice had greater modularity than water drinking female controls [41]. Similarly, chronic binge-like ethanol drinking increased modularity relative to water drinkers in socially housed female mice, and re-exposure to ethanol or ethanol-paired cues further increased brain network modularity [40]. Following 8 days of binge-like drinking, Ardinger et al., also found that female ethanol drinking mice have greater modularity than water drinking counterparts. Similarly to Kimbrough et al. and Bloch et al., this study found that male ethanol drinking mice had lower modularity than male water controls [38]. Overall, the present study agrees with the 2 previous reports that ethanol drinking increases modularity in female mice, regardless of the number of drinking days or drinking model. However, our study found that male ethanol drinking mice had greater modularity than male water drinking mice. These differences could be attributed to the drinking model, history of drinking (4 versus 8+ days), physiological state (intoxicated versus in withdrawal or abstinence), or time of euthanasia [time of day and time since start of drinking (4hr versus 30-80 minutes)]. Modularity is an important metric that allows us to evaluate what regions are most closely connected to each other, and how ethanol perturbs these connections. Together, results indicate ethanol increased modularity in females, but effects of ethanol on modularity of male networks depends on the drinking model.

### Evidence of hemisphere differences

Assessment of hemisphere differences have often been neglected in the rodent literature, likely due to technical limitations. There is evidence of lateralized structure and function of the rodent amygdala [56–60]. The present results have also identified hemisphere differences in amygdalar nuclei that are NAcc inputs differentially engaged by ethanol drinking (left anterior amygdala) and module hubs (right intercalated amygdala, left cortical and basomedial amygdala). Imaging studies in humans, non-human primates, and rodents have identified alcohol-related changes in neural responses, connectivity, and network dynamics that are often biased to one hemisphere [61–65]. Of the 129 regions with greater c-Fos in female relative to male ethanol drinkers, there was significant overrepresentation of regions in the right hemisphere (89 right versus 40 left). However, this was not true of NAcc inputs with greater engagement in female relative to male ethanol drinkers (22 right, 16 left). An important implication of these findings is that future studies should consider hemisphere differences and lateralization of function when technically feasible.

### Limitations

c-Fos is only one immediate early gene and does not capture the full temporal scope of region activation. Future studies should consider using other immediate early genes, either alone or in combination with c-Fos. Arc may more sensitively capture neuronal activity levels [66], and the recently described marker of inhibition, phosphorylation of pyruvate dehydrogenase, could disentangle the pharmacological inhibitory from the stimulatory effects of ethanol [67], but both would require more sophisticated automated cell detection algorithms for whole brain studies.

Another consideration is whether c-Fos induction is entirely neuron specific. Evidence from *in vitro* studies found that ethanol delayed the downregulation of c-Fos expression in oligodendrocyte precursor cells [68], and induced c-Fos expression in astrocytes [69]. Although our tissue was labeled with NeuN for image alignment, differential NeuN staining based on tissue depth prevents reporting of colocalization.

Use of the B6 mouse has limitations, including a genetic mutation that reduces expression of the alpha-2 subunit of the GABA-A receptor, which likely impacts the pharmacological effect of alcohol [70–72]. Additionally, B6 is an inbred mouse strain, and does not capture the polygenicity of alcohol drinking and AUD in humans [73]. Future studies could employ genetic models of risk for drinking to intoxication (e.g. High Drinking-in-the-Dark mice) to assess the contribution of genetic risk on brain region engagement [74–76].

### Conclusions & Future Directions

We identified several regions and NAcc inputs differently engaged by ethanol, as well as novel hub regions with unique connectivity in male and female networks. Far more regions and NAcc inputs were engaged by binge-like ethanol drinking in female than male mice. However, binge-like ethanol drinking increased network modularity and decreased network connectivity more dramatically in males. Together, these findings further our understanding of how ethanol significantly and differentially effects the male and female mouse brain.

## Acknowledgements

Thank you to Joselinne Medrano, Zaynah Usmani, and Courtney Ledford for technical assistance. Thank you to members of the Ozburn (Louis Nuñez, Luis Tzab, Ian Anderson) and Ryabinin (Michael Johnson, Kelly Abshire, Jonathan Zweig) labs for helpful discussions during preparation of this manuscript. We also acknowledge the expert technical assistance by the OHSU Advanced Light Microscopy Core in 3D visualization of the cleared mouse brain (RRID:SCR_009961). This work was supported by the US Department of Veterans Affairs Merit Award (I01 BX0046990, I01BX006570), National Institute on Alcohol Abuse and Alcoholism (AA013519, AA010760, AA030908, AA007468, AA030806, AA028680), National Institute on Drug Abuse (R25 DA050727), ARCS Foundation-Oregon Chapter, and a generous gift from the John R. Andrews family.

## Author Contributions

AEC performed the experiments and analyses, interpreted the analysis, and wrote the paper. JQA performed and interpreted the statistical analysis and edited the paper. KBG and BEJ assisted with performing the experiments and edited the paper. AER and ARO interpreted results and conclusions. ARO conceived the experiment, planned and interpreted analysis, and wrote the paper.

## STAR Methods

### Resource availability

#### Lead contact

Further information should be directed to and fulfilled by the lead contact, Dr. Angela Ozburn

#### Materials availability

This study did not generate unique reagents.

### Data and code availability

Datasets will be deposited at the time of publication. Image files will be deposited at the Brain Image Library at the time of publication (https://www.brainimagelibrary.org/).

Source Code will be deposited at GitHub at the time of publication (https://github.com/OzburnLabOHSU/NAcc-Circuitry---Chan-et-al.-2024).

Any additional information required to reanalyze the data reported in this paper is available from the lead contact upon request.

### Experimental model details

#### Mice

Adult male and female C57BL/6J (B6, Cat#000664; RRID: MSR_JAX:000664; 10 weeks old, 24.4g ± 0.49 at start of experiment) mice were obtained from Jackson Laboratory (Sacramento, CA) and housed at 22 ± 1^°^C on a 12hr/12hr reverse light/dark cycle with lights off at 0730 (PST). The B6 mouse was chosen because they reliably drink to intoxication in the Drinking in the Dark (DID) paradigm, including studies that found sex differences in ethanol intake when NAcc activity was chemogenetically manipulated [19–21]. Mice were acclimated to this reverse light/dark cycle for 1 week prior to surgery. Mice had access to tap water and standard chow (5LOD, PMI Nutrition International, St. Louis, MO) *ad libitum* unless otherwise noted. Mice were group housed until 1 week prior to the start of DID, when they were singly housed for the rest of the experiment. All protocols were approved by the Portland Veteran Affairs Medical Center Institutional Animal Care and Use Committees and were conducted in accordance with the National Institutes of Health Guidelines for the Care and Use of Laboratory Animals.

### Method details

#### Stereotactic Surgery

All mice underwent stereotactic surgery under isoflurane anesthesia (5% induction, 1-3% maintenance). Intracranial viral injections were carried out using standard procedures [19, 52, 77]. AAVrg-hSyn-eGFP (0.5 µl/side, Addgene, Watertown MA), was injected bilaterally into the NAcc (10° angle, M/L ± 1.50, A/P + 1.50, D/V −4.50 and −4.00 mm from bregma) using a 33-gauge Hamilton syringe (Hamilton, Reno NV). Virus was infused at a rate of 0.1µl/minute, and the needle was kept in place for an additional 10 minutes to allow proper diffusion. 0.25µL of virus was delivered at −4.50mm, and the remaining 0.25µL of virus was infused at −4.00mm. pAAV-hSyn-eGFP was a gift from Bryan Roth (Addgene viral prep Cat#50465-AAV-rg). For details on the development and validation of this viral construct see [78]. Following surgery, mice were given ketofen (5mg/kg, s.c.) and saline (0.5mL, s.c.) daily for 3 days. Experiments began 3 weeks after surgery to allow for full viral expression (See **Figure 1A** for experimental timeline).

#### Drinking-in-The-Dark

Drinking in the Dark (DID) is a limited access drinking assay that consistently results in B6 mice achieving intoxicating blood ethanol concentrations (BEC; [32]). Briefly, 3h into the dark cycle, water is removed from the home cage and replaced with a volumetric sipper containing 20% ethanol (v/v in tap water) or water. On days 1-3 of DID, mice have measured fluid access for 2hrs, and on day 4, mice have measured fluid access for 4hrs. After the access period, sippers are removed, and water bottles are returned. Immediately after the 4hr DID session, 20µL of periorbital sinus blood was collected from ethanol drinking mice. BECs were determined using gas chromatography as described in [79]. Water drinking mice were handled in a similar manner, except no blood was collected.

#### Tissue Collection

Immediately following blood collection, mice were deeply anesthetized with a cocktail of ketamine (200mg/kg) and xylazine (20mg/kg) prior intracardial perfusion for 5 minutes with phosphate buffered saline (PBS) containing 10U/mL heparin, followed by 4% paraformaldehyde (PFA) in PBS. Brains were removed and post-fixed overnight. Brains were transferred to PBS with 0.02% sodium azide until tissue clearing.

#### Tissue Processing

Tissue processing was conducted by LifeCanvas Technologies (LCT, Cambridge MA) following their established SHIELD protocol [80]. Samples were SHIELD post-fixed at LCT, followed by 7-days of delipidation in Clear+. Samples were labeled for c-Fos (Rabbit anti-cFos, Abcam Cat#214672; RRID:AB_2939046, 3.5µg/batch), GFP (goat anti-GFP, Encor Cat#GPCA-GFP; RRID:AB_2737371, 5µg/batch), and NeuN (mouse anti-NeuN, Encor Cat#MCA-1B7; RRID:AB_2572267, 12µg/batch). Samples were delipidated and antibody labeled using the SmartBatch+ device in batches of 12 brains [81]. Images were acquired on the SmartSPIM light-sheet microscope with a voxel resolution of 1.8μm x 1.8μm x 4μm (z-step).

#### Image visualizations

SmartAnalytics (LifeCanvas Technologies) was used to produce heatmaps of c-Fos density from representative mice (**Figure S2**). ImageJ (NIH) was used to visualize GFP expression [82]. Imaris (Bitplane, version 10.1.0) was used to pseudo-color immunolabelled cells, render 3D video, and visualize example c-Fos+GFP colocalization from a representative male, ethanol drinking mouse (**Figure S5** and **Video S1**).

#### Viral placement

Only mice with bilateral NAcc GFP expression were included in the c-Fos+GFP colocalization analysis (n=7-14/sex/fluid). This subset of mice had the same drinking behavior and BECs as the larger sample (**Figure S4A-E**). Mice were considered bilateral hits if the majority (>50%) of viral expression was in the NAcc, though there was some viral spread in the medial nucleus accumbens shell. All mice had similar anterior-posterior viral spread (**Figure S4F**). Mice with unilateral GFP expression were excluded because the bilateral viral injections make identification of ipsilateral versus contralateral NAcc inputs beyond the scope of this project. Variability in the spread of the viral tracer prevents us from quantifying the proportion of NAcc inputs that are engaged.

### Quantification and statistical analysis

#### Image processing & cell quantification

Images were registered to the Allen Brain Institute Common Coordinate Framework using a two-phase semi-automated process in the SmartAnalytics software [83–85]. In the automated phase, NeuN channel images of each brain were registered to atlas-aligned reference samples, using successive rigid, affine, and b-spline warping algorithms (SimpleElastix [86]). An average alignment to the atlas was created across all reference sample alignments. During the manual phase, each alignment was viewed with a custom Neuroglancer interface and correspondence points between the atlas and the sample are adjusted if needed. Next, automated cell counts of c-Fos, GFP, co-labeled c-Fos and GFP positive cells was performed using a custom convolutional neural network created by LCT with the Tensorflow Python package [87, 88]. To detect cells in 3D microscope images, a two-step approach consisting of a candidate detection network and a cell classification network was employed. The candidate detection network aimed to identify all possible cell locations while minimizing false negatives. For this purpose, we utilized a U-Net, a fully convolutional network known for its effectiveness in semantic segmentation tasks. The candidate detection network was trained using the hand-tagged ROI where positive cell locations were labeled. The goal of the cell classification network was to classify the locations identified by the candidate detection network as either positive or negative cells. To maximize total accuracy a 3D ResNet was used as the structure for the classification convolutional neural network. Training data for the cell classification network consisted of small regions surrounding cells identified by the candidate detection network on the training regions of interest. Cells positively labeled by a researcher served as positive training examples, and candidate regions that did not overlap with true cell locations served as negative training examples.

#### Drinking data and BEC data analysis

Data are shown as mean ±SEM. Data were analyzed using a 2-way ANOVA with 2hr fluid intake as repeated measures (2hr intake days 1-4) and Student’s unpaired two-tailed t-test (day 4 4hr intake and BECs). Nearly all mice drank to intoxication, so we did not correlate BEC with c-Fos expression. All main effects and interactions are reported, with post hoc testing using Šidák’s multiple comparisons test when appropriate. An alpha level of 0.05 was used to determine significance. Data analysis was performed, and graphs were generated using GraphPad Prism 10.

#### Preliminary image data analysis

RStudio (version 06.2.561) and the tidyverse package (version 2.0.0 [89]) were used for analysis of whole-brain c-Fos expression and c-Fos+GFP colocalization and for the generation of **Figures 2-4**. The raw output of the cell detection algorithm contains data (cell density, cell/mm^3^) for 1678 Allen Brain Atlas defined regions per mouse, including layer specific information from the cortex, white matter, ventricles, hindbrain structures, and separate values for left and right hemispheres. This data was reduced to 426 areas per mouse (213 per hemisphere), eliminating values for white matter and ventricles, and reducing subregion and layer specific information (ex. infralimbic cortex layer I, II/III, V, VIa, and VIb were collapsed into 1 value for this region, **Table S1**).

#### Data Distribution

The distribution of whole-brain density data was visualized through violin plots using the *ggstatsplot* package (version 0.11.1 [90]). Visualization of un-transformed c-Fos density and c-Fos+GFP colocalization data indicated a rightward skew with no statistical outliers. All c-Fos density and c-Fos+GFP colocalization data was square root transformed to approximate a normal distribution (**Figures S1 and S6,** [91]). Reduced, square root normalized density data was used for all further analyses of c-Fos and c-Fos+GFP colocalization.

To determine whether there were overall differences in c-Fos expression, a 1-way ANOVA was used to compare whether total c-Fos density per mouse differs between sex and fluid groups. There was no effect of sex and fluid groups on c-Fos density [F_(3,57)_=1.87, p>0.05]. Likewise, there were no differences between sex and fluid groups in total c-Fos+GFP colocalization [F_(3,37)_=0.475, p>0.05]. To determine whether there were differences in c-Fos density between hemispheres, c-Fos density was summed per hemisphere (2 values per mouse) and sex and fluid groups were compared with a Welch’s paired t-test. There was no effect of hemisphere on total c-Fos density, and similarly no effect of hemisphere on c-Fos+GFP colocalization.

#### Principal component analysis (PCA)

Dimensionality reduction was performed using the *prcomp* function of R’s *stats* package (version 4.1.2) to identify the major sources of variance in the data (c-Fos see **Figure 2** and **Table S2**, c-Fos+GFP see **Figure S6** and **Table S4**). The top 10 PCs (explaining ∼72% of variance) were included in analysis. Association between PCs and factors were determined by 1-way ANOVA (significance threshold of p<0.05). Two-way ANOVA [*anova_test* function of *rstatix* package (version 0.7.2)] was used to determine whether sex by fluid interaction was significantly associated with any of the top 10 PCs.

#### Comparison of c-Fos density profile

To visualize c-Fos expression patterns between sex and fluid groups, hierarchically clustered heatmaps were created with the *ComplexHeatmap* function *Heatmap* (version 2.10.0 [92, 93]). Average c-Fos density values for each region are plotted by sex and fluid groups. The complete clustering algorithm was used to cluster regions (rows) by similarity in c-Fos density, and sex and fluid groups (columns) by similarity of c-Fos density profile (**Figure 2A**).

#### Differential expression

Visualization of regions with different c-Fos density or c-Fos+GFP colocalization across conditions was done using volcano plots created with ggpubr (version 0.6.0 [94]). Log-fold change was calculated between two conditions for each region [ex: log_2_(mean square root transformed c-Fos density in region A of ethanol drinking mice) - log_2_ (mean square root transformed c-Fos density in region A of water drinking mice)] and plotted on the x-axis. A Welch’s t-test was used to calculate a p-value for each region to compare c-Fos density or c-Fos+GFP colocalization between two conditions (**Figure 2D-G, S3, S7, Tables S3, S5**).

### c-Fos Expression Network construction

#### Method 1: Correlation Matrix Networks

A correlation matrix was created for each sex and fluid group by calculating Pearson’s R values for each pair of regions using c-Fos density (**Figure 3A-D, Tables S7, S9, S11, S13**). This method was chosen based on prior work [38–41] and has been employed to successfully identify a region of interest causally linked to dependence-like ethanol intake in males [55]. Hierarchical clustering was used to group regions using the average method (*hclust, stats* package) and cutting the clustering dendrogram at a set height (*cuttree, stats* package) was used to determine modules. A tree cut height of 70% was chosen because it retained group differences in number of modules that existed at tree cut heights 90% and below. Correlation heatmap matrices were created with *corrplot* (Version 0.92 [95]).

#### Method 2: WGCNA

For the Weighted Gene Co-expression Network Analysis (WGCNA) approach, we created an overall network using c-Fos density from all sex and fluid groups. This method constructs a scale-free network that models the structure of biological networks (for review on brain network structure see [54]). The eigengene networks constructed with WGCNA can identify biologically meaningful networks from gene transcription data [96]. Data from one male water drinking mouse was determined to be an outlier and excluded from analysis (n=14-16/sex/fluid). Using the *blockwiseModules* function of the *WGCNA* package ([97] version 1.72-1), we created an unsigned network with the following parameters: power = 8, minModuleSize = 10, mergeCutHeight = 0.15, pamRespectsDendro = FALSE, and deepSplit = 2. Thes parameters constructed a network of 17 modules (**Figure 4A, Table S16**). To determine whether any modules were associated with categorical variable(s), a 2-way ANOVA (factors: sex and fluid) was completed using module eigenregions (MEs). To determine whether the MEs from any module in ethanol drinking mice are significantly correlated with ethanol consumption (BEC, 4hr ethanol intake, or total 4-day ethanol intake), Pearson’s correlations values were calculated using the *cor* (*stats* package), and p-values were calculated with the *corPvalueStudent* (*WGCNA* package).

WGCNA identified modules associated with total 4-day ethanol intake that were not associated with day 4, 4hr ethanol intake (green, grey, light cyan, pink, and purple) and vice versa (turquoise, **Figure 4B, Table S16**). Four modules (grey, light cyan, pink, purple) are associated with both total 4-day ethanol intake and sex, so regions in these modules are likely influenced by the different levels of total ethanol intake between males and females.

#### Network module characterization

For each network, the proportion of regions per anatomical category within each module were plotted (**Figures S8A**, **S10A, Tables S6, S8, S10, S12**). Anatomical groups (cerebellum, cortical plate, cortical subplate, hindbrain, hypothalamus, midbrain, pallidum, striatum, and thalamus) were defined using the Allen Brain Atlas [85] and visualized using the *ggplot2* package using perceptually uniform color maps (*viridis* package, version 0.6.3 [98]). Fisher’s exact tests (*stats* package) were used to determine whether any anatomical category was significantly overrepresented [false discovery rate (fdr) corrected p<0.05] in a module. Similarly, we plotted and determined whether regions from the left or right hemisphere were overrepresented in the modules of each network (**Figures S8B, S10B, Tables S6, S8, S10, S12**). More modules in the WGCNA network had significant overrepresentation of a hemisphere (fdr’s<0.05) than modules in the correlation matrix networks.

For the correlation matrix networks, plots of the average correlation coefficients of each region within a module was created with *ggstatsplot* function *ggbetweenstats* (**Figure S8C**). The largest modules in these networks tended to have the highest average intramodular correlation coefficients. A comparison of average correlation coefficients for regions within each anatomical group between each sex and fluid group was also calculated (**Figures S9A-H**). Significant differences between groups were determined by Welch’s t-test and Holm’s adjusted p-values.

For the WGCNA network, a plot of the average adjacency value for each region in a module was created with *ggstatsplot* function *ggbetweenstats* (**Figure S10C**). Significant differences between modules were determined by Welch’s t-test and Holm’s adjusted p-values.

#### Module Preservation

Chord diagrams were created to visualize the overlap of regions within network modules across correlation matrix networks using the *ChordDiagram* function of *circilize* package (version 0.4.15 [99]). Fisher’s exact tests (*stats* package) were used to determine whether there was significant overrepresentation of regions between modules of different sex and fluid group networks (**Figures 3E-H**, **Tables S6, S8, S10, S12**, comparisons made across sex or fluid).

#### Differential Wiring

To determine how ethanol changes regional connectivity, we identified regions pairs with significant changes in correlation values between same-sex water and ethanol networks (**TablesS S14, S15**). The male water was compared to the male ethanol network, and the female water was compared to the female ethanol network. Threshold for change in correlation was fdr<0.05, ΔR≥0.5 with at least 1 other region. Log fold change was calculated as follows: log_2_(abs(sum correlations region A in ethanol network))/(abs(sum correlations of region A in water network)).

#### Anti-correlations

Prior work found that increased anti-correlations are associated with lower activity of parvalbumin inhibitory interneurons [100, 101]. To determine whether sex or fluid influence the proportions of negative and positive correlations in a network, we counted of the number of negative and positive correlations with the *signcount* function of the *wordspace* package (version 0.2-8 [102]). Fisher’s exact tests (*stats* package) were used to determine whether there was a significant difference in number of negative and positive correlations within sex and fluid groups.

The male water network had a significantly lower percentage of negative correlations (3.17%), sex (male ethanol, 20.6%,) and fluid matched (female water, 17.6%) networks. The female water network a significantly lower percentage of negative correlations the female ethanol (204.4%) network. The female and male ethanol networks had comparable percentages of negative correlations (20.4% and 20.6%, respectively). This suggests that changes in network connectivity are due to greater reduction in parvalbumin inhibitory interneuron activity in male than female ethanol drinkers.

#### Network Topology

The participation coefficient (P) is a ratio of the intra-versus inter-module connections of each region in a network. P was calculated using values from the *WGCNA* package function *intramodularConnectivity* as follows: 1-(kWithin/kTotal) where kWithin is a region’s total intramodular connectivity and kTotal is a region’s total connectivity. Non-hub nodes were classified as ultra-peripheral (P≤0.05, only intramodular connections), peripheral (0.05< P ≤0.62, mostly intramodular connections), non-hub connector (0.62<P≤0.80, mostly inter-module connections), or non-hub kinless (P>0.80, evenly connected with all modules) as defined in [42]. Hubs, defined as regions with the top 20% of intramodular connectivity, were classified as provincial (P<0.3, mostly intramodular connections), connector (0.3<P<0.75, mostly inter-module connections), or kinless (P>0.75, connections distributed across all modules). Plots of the number of each type of node and hub were created with the base R function *pie* and the *RColorBrewer* package (version 1.1-3 [103], **Figures S8D, S10D, Tables S7, S9, S11, S13, S17).**

Network graphs were created using the *igraph* (version 1.4.2 [104, 105]), *tidygraph* (version 1.2.3 [106]), and *ggraph* (version 2.1.0 [107]) packages. Graphs were created from adjacency matrices for regions in the WGCNA network midnight blue module for each sex and fluid group separately (**Figure 4C**). The Fruchterman-Reingold (fr) algorithm was used to plot the graphs. Node size denotes importance of node in network (larger nodes are hubs). Opacity and width of arcs denotes strength of connections.

## Supplemental Information

**Supplemental Figure S1.** Reduced untransformed and transformed data distribution (A) Violin plots of raw, reduced c-Fos density to visualize data distribution. (n=15-16/sex/fluid, 426 data points per mouse). (B) Violin plots or square root (sqrt) transformed, reduced c-Fos density to visualize data distribution. (C) Sum of c-Fos density (reduced, sqrt transformed) in each sex and fluid group. No difference in c-Fos density between groups [1-way ANOVA, p>0.05, F_(3,57)_=1.87]. (D) Sum of c-Fos density (reduced, sqrt transformed) per hemisphere (2 values per mouse), sex and fluid groups collapsed. No difference in c-Fos density between groups (Welch’s paired t-test, p>0.05).

***Supplemental Figure S2:*** *Representative heatmaps of c-Fos expression* (A-D) Coronal c-Fos density (cells/mm^3^) heatmaps from male water (A), female water (B), male ethanol (C), and female ethanol (D) drinking mice. Images created with SmartAnalytics. Coordinates given in mm from bregma. NAcc, nucleus accumbens core; PVN, periventricular nucleus of the hypothalamus; VP, ventral posterior thalamus.

***Supplemental Figure S3.*** *Differential c-Fos expression* Volcano plots comparing c-Fos density between different conditions. Dotted line at –log_10_(p-value)=1.3 denotes threshold for significance. (A) 11 regions had significantly higher c-Fos expression in water than ethanol drinking mice (n=30-31/fluid, sex collapsed). 6 regions had significantly higher c-Fos expression in ethanol than water drinking mice. (B) No regions had significantly greater c-Fos expression in male than female mice (n=30-31/sex, fluid collapsed). 90 regions had significantly greater c-Fos expression in female than male mice. (C) 2 regions had significantly greater c-Fos expression in female water than female ethanol drinking mice (n=15-16/fluid). 15 regions had significantly greater c-Fos expression in female ethanol than female water drinking mice. (D) 12 regions had significantly greater c-Fos expression in male water than male ethanol drinking mice (n=15/fluid). 1 region had significantly greater c-Fos expression in male ethanol than male water drinking mice. See **Table S3** for additional information.

***Supplemental Figure S4.*** *Drinking data for mice with bilateral NAcc hits* (n=7-14/sex/fluid). (A) 2hr ethanol intake over 4-day DID, significant effect of sex (n=10-14/sex, day 1-3 and day 4 0-2hr intake, F_(1,22)_= 26.25; ****p<0.0001), no effect of day, or sex by day interaction. (B) Day 4 4hr ethanol intake. No effect of sex. (C) BEC after 4hr ethanol intake. No effect of sex. (D) 2hr water intake over 4-day DID (n=7-10/sex). No effect of sex, day, or sex by day interaction. (E) Day 4 4hr water intake. No effect of sex. Closed circles denote males, open triangles denote females. Data shown as mean ± SEM. (F) Illustration of maximal GFP viral expression in the NAcc. Images adapted from *The Mouse Brain in Stereotaxic Coordinates*, Franklin & Paxinos (2007). Pink shapes denote females, orange shapes denote males.

***Supplemental Figure S5.*** *Representative c-Fos+GFP colocalization and expression* Representative c-Fos and GFP expression from a male ethanol drinking mouse. (A) Expression of c-Fos (magenta) and GFP (cyan) in right hemisphere medial prefrontal cortex. Scale bar = 300um. (B) Higher magnification image of region outlined in white box (infralimbic cortex) illustrating c-Fos expression (top), GFP expression (middle), and their overlap (bottom). Arrows in bottom panel indicate white, co-labeled c-Fos+GFP cells. Scale bar = 300um. (C) Whole brain image of c-Fos and GFP expression. Scale bar = 800um. A, anterior; D, dorsal; P, posterior; V, ventral.

***Supplemental Figure S6.*** *c-Fos+GFP reduced data distribution and Principal Component Analysis* (A) Violin plots of raw, reduced c-Fos+GFP colocalization density (cells/mm^3^) to visualize data distribution (n=7-14/sex/fluid, 423 data points per mouse). (B) Violin plots of square root (sqrt) normalized, reduced c-Fos+GFP colocalization to visualize data distribution. (C) Sum of c-Fos+GFP colocalization (reduced, sqrt transformed) in each sex and fluid group. No difference in total colocalization between groups [1-way ANOVA, p>0.05, F_(3,37)_=0.475]. (D) Sum of c-Fos+GFP colocalization (reduced, sqrt transformed) per hemisphere (2 values per mouse), sex and fluid groups collapsed. No difference in total colocalization between hemispheres (Welch’s paired t-test, p>0.05). (E&F) Top 2 PCs (determined by 1-way ANOVA) and proportion of explained variance for each factor of interest for normalized, reduced c-Fos+GFP colocalization. (E) Fluid is significantly associated with PC5 (p<0.05, 4.8% of total variance). (F) Sex is significantly associated with PC7 (p<0.05, 3.7% of total variance) and PC10 (p<0.05, 2.3% of total variance). See **Table S4** for additional information.

***Supplemental Figure S7.*** *Differential c-Fos+GFP colocalization* Volcano plots comparing c-Fos+GFP colocalization between different conditions. Dotted line at – log_10_(p-value)=1.3 denotes threshold for significance. (A) 4 regions had significantly higher c-Fos+GFP colocalization in water than ethanol drinking mice (n=17-24/fluid, sex collapsed). 10 regions had significantly higher c-Fos+GFP colocalization in ethanol than water drinking mice. (B) 1 region had significantly greater c-Fos+GFP colocalization in male than female mice (n=17-24/sex, fluid collapsed). 36 regions had significantly greater c-Fos+GFP colocalization in female than male mice. (C) 2 regions had significantly greater c-Fos+GFP colocalization in female water than female ethanol drinking mice (n=10-14/fluid). 9 regions had significantly greater c-Fos+GFP colocalization in female ethanol than female water drinking mice. (D) *3* regions had significantly greater c-Fos+GFP colocalization in male water than male ethanol drinking mice (n=7-10/fluid). 1 region had significantly greater c-Fos+GFP colocalization in male ethanol than male water drinking mice. See **Table S5** for additional information.

***Supplemental Figure S8.*** *Correlation matrix network module characterization* Clusters determined from hierarchically clustered Pearson’s correlation matrix at 70% tree cut height for each sex and fluid group (n=15-16/sex/fluid). (A) Proportion of regions per anatomical category in each module. (B) Proportion of regions per hemisphere in each module. (C) Average intramodular correlation coefficient for each region within a module. Welch’s t-test with Holm’s multiple comparison’s corrected p-values. Created with ggbetweenstats from ggpubr. *p< 0.05, **p< 0.01, ***p< 0.001, ****p< 0.00001. (D) Proportion of regions in each node and hub classification. See **Tables S6, S8, S10, S12** for additional information.

***Supplemental Figure S9.*** *Average correlation coefficient per anatomical subgroup and largest module for each correlation matrix network.* (A-H) Anatomical divisions were defined by the Allen Brain Atlas (n=15-16/sex/fluid). Welch’s t-test with Holm’s multiple comparison’s corrected p-values. Main effect of group in cortical plate (F_(3,270.81)_=174.92, p<0.00001), cortical sublate (F_(3,24.14)_=65.93, p<0.00001), hindbrain (F_(3,90.86)_=75.61, p<0.00001), hypothalamus (F_(3,166.46)_=17.44, p<0.00001), midbrain (F_(3,144.02)_=51.41, p<0.00001), pallidum (F_(3,37.38)_=58.92, p<0.00001), striatum (F_(3,41.92)_=45.87, p<0.00001), and thalamus (F_(3,143.27)_=248.09, p<0.00001). Created with ggbetweenstats from ggpubr. *p< 0.05, **p< 0.01, ***p< 0.001, ****p< 0.00001. Cerebellum excluded because there is only 1 data point per group (correlation between right and left cerebellum). (I) Average intramodular correlation value for each region within the largest network module of each sex and fluid group. Main effect of group (F_(3,552.57)_=184.21, p<0.00001). Significant differences determined by Welch’s t-test, Holm’s adjust p-values reported. ****p< 0.00001.

***Supplemental Figure S10.*** *WGCNA network characteristics* (A) Proportion of regions per anatomical category in each module. (B) Proportion of regions per hemisphere in each module. (C) Adjacency values per module. Significant effect of module [F_(16,113.68)_=61.56, p<0.00001]. Table to the right contains significant module pair-wise comparisons of adjacency values. *D.* Proportion of regions in each node and hub classification.

