## Supplemental Figures for "Sex differences in nucleus accumbens core circuitry engaged by binge-like ethanol drinking"

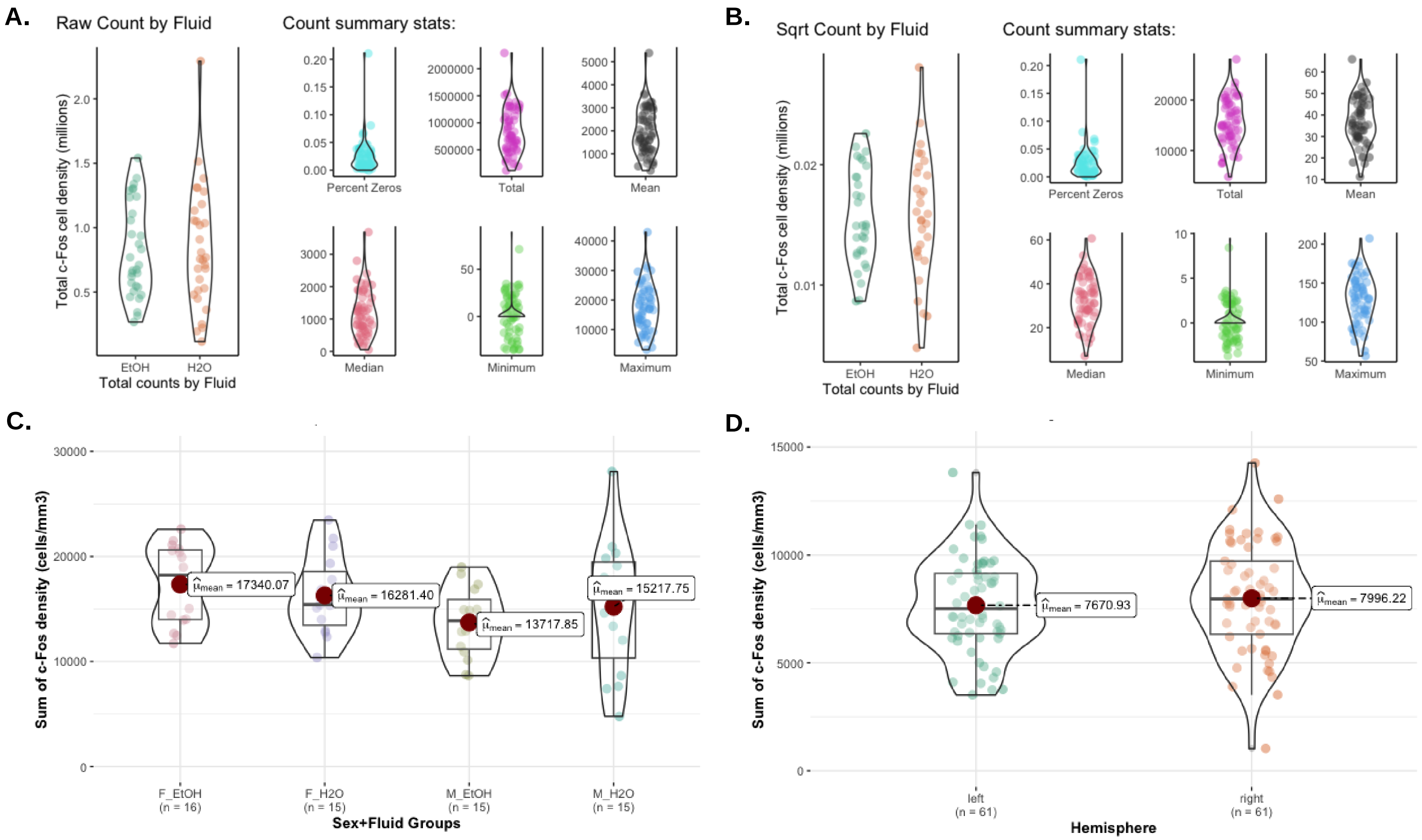
A.
Raw Count by Fluid
Count summary stats:
Total c-Fos cell density (millions)
Total counts by Fluid
EtOH
H2O
Percent Zeros
Total
Mean
Median
Minimum
Maximum
B.
Sqrt Count by Fluid
Count summary stats:
C.
Sum of c-Fos density (cells/mm3)
$\hat{\mu}_{mean}$ = 17340.07
$\hat{\mu}_{mean}$ = 16281.40
$\hat{\mu}_{mean}$ = 13717.85
$\hat{\mu}_{mean}$ = 15217.75
F_EtOH (n = 16)
F_H2O (n = 15)
M_EtOH (n = 15)
M_H2O (n = 15)
Sex+Fluid Groups
D.
$\hat{\mu}_{mean}$ = 7670.93
$\hat{\mu}_{mean}$ = 7996.22
left (n = 61)
right (n = 61)
Hemisphere

Figure S1

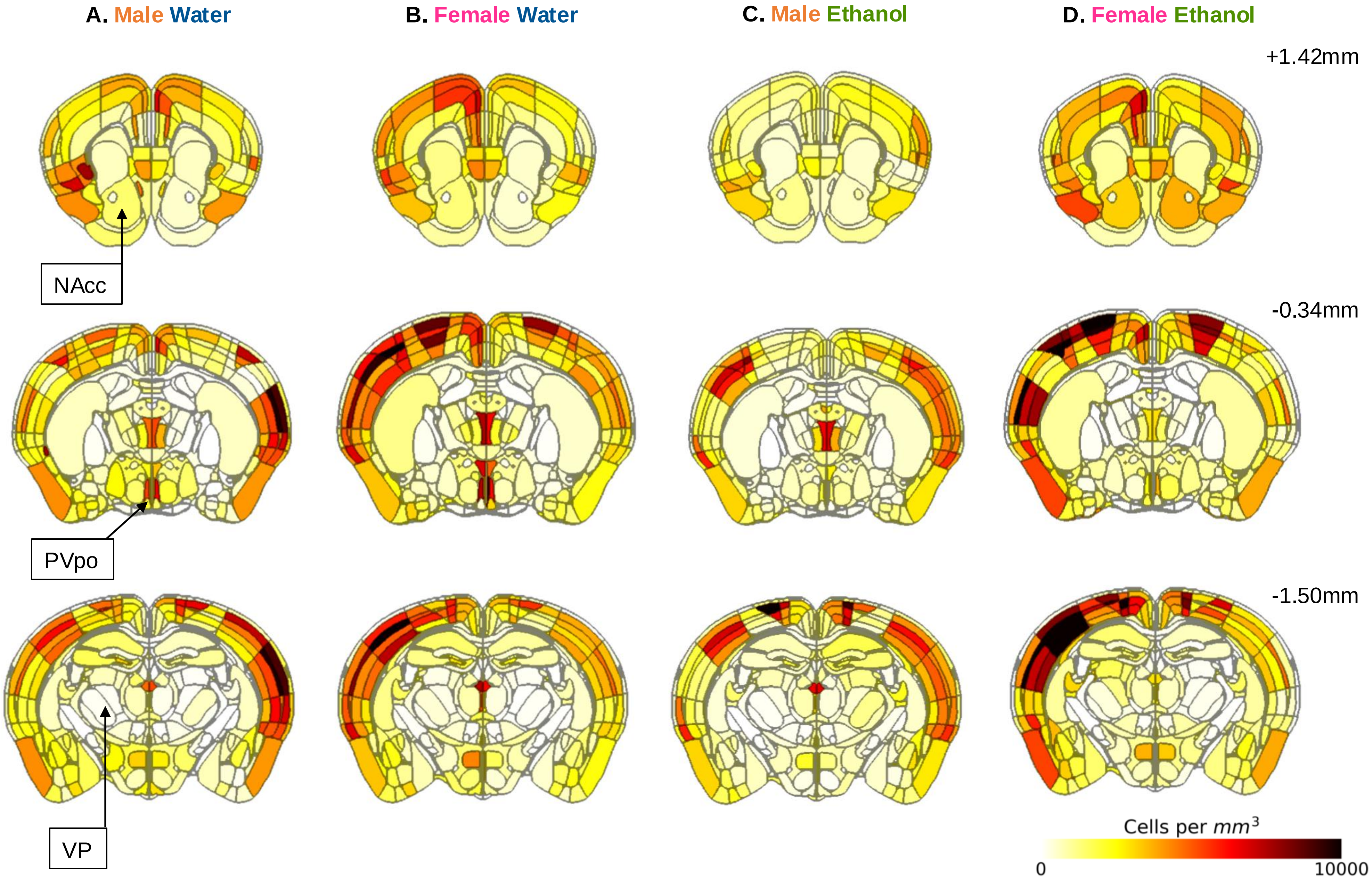
A. Male Water
B. Female Water
C. Male Ethanol
D. Female Ethanol
+1.42mm
-0.34mm
-1.50mm
NAcc
PVpo
VP
Cells per $mm^3$
0
10000

Figure S2

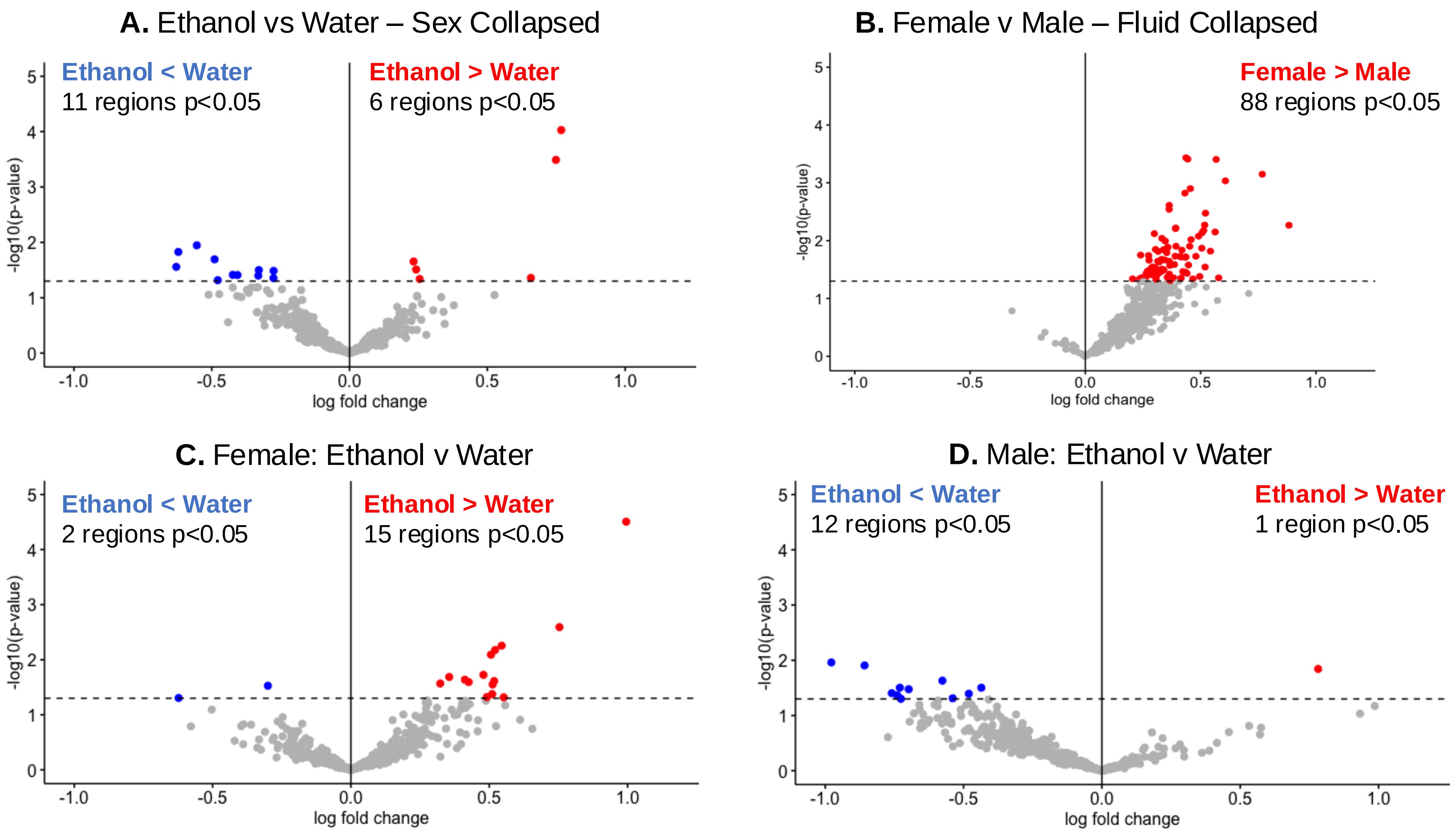

A. Ethanol vs Water – Sex Collapsed
Ethanol < Water
11 regions p<0.05
Ethanol > Water
6 regions p<0.05
-log10(p-value)
log fold change
B. Female v Male – Fluid Collapsed
Female > Male
88 regions p<0.05
-log10(p-value)
log fold change
C. Female: Ethanol v Water
Ethanol < Water
2 regions p<0.05
Ethanol > Water
15 regions p<0.05
-log10(p-value)
log fold change
D. Male: Ethanol v Water
Ethanol < Water
12 regions p<0.05
Ethanol > Water
1 region p<0.05
-log10(p-value)
log fold change


Figure S3

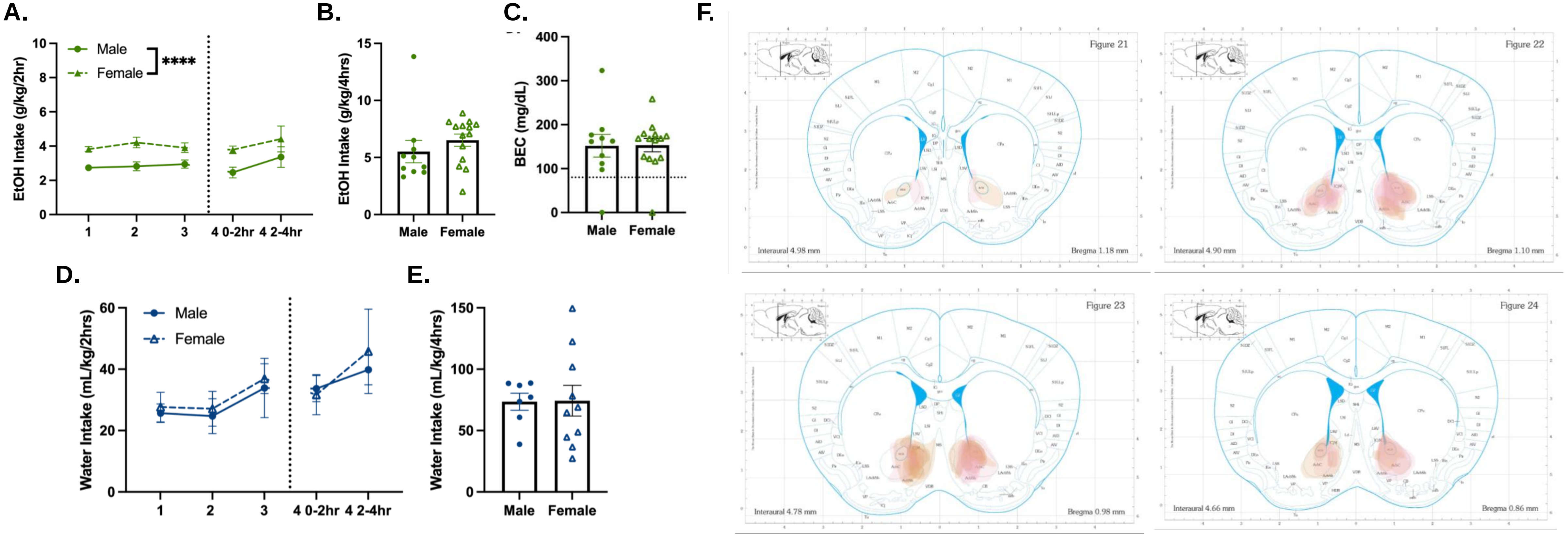
A.
EtOH Intake (g/kg/2hr)
Male
Female
****
1
2
3
4 0-2hr
4 2-4hr
B.
EtOH Intake (g/kg/4hrs)
Male
Female
C.
BEC (mg/dL)
Male
Female
D.
Water Intake (mL/kg/2hrs)
Male
Female
E.
Water Intake (mL/kg/4hrs)
Male
Female
F.
Figure 21
Figure 22
Figure 23
Figure 24
Interaural 4.98 mm
Bregma 1.18 mm
Interaural 4.90 mm
Bregma 1.10 mm
Interaural 4.78 mm
Bregma 0.98 mm
Interaural 4.66 mm
Bregma 0.86 mm

Figure S4

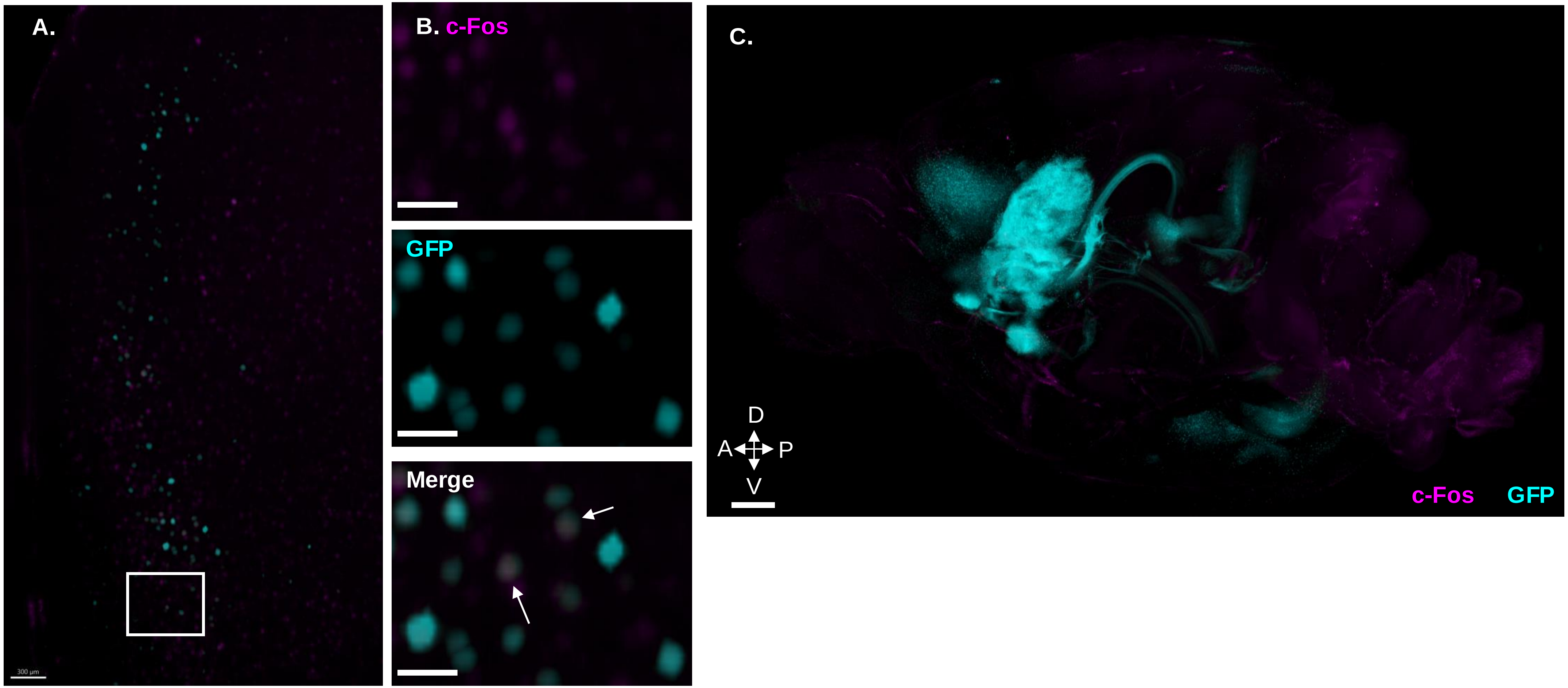


Figure S5

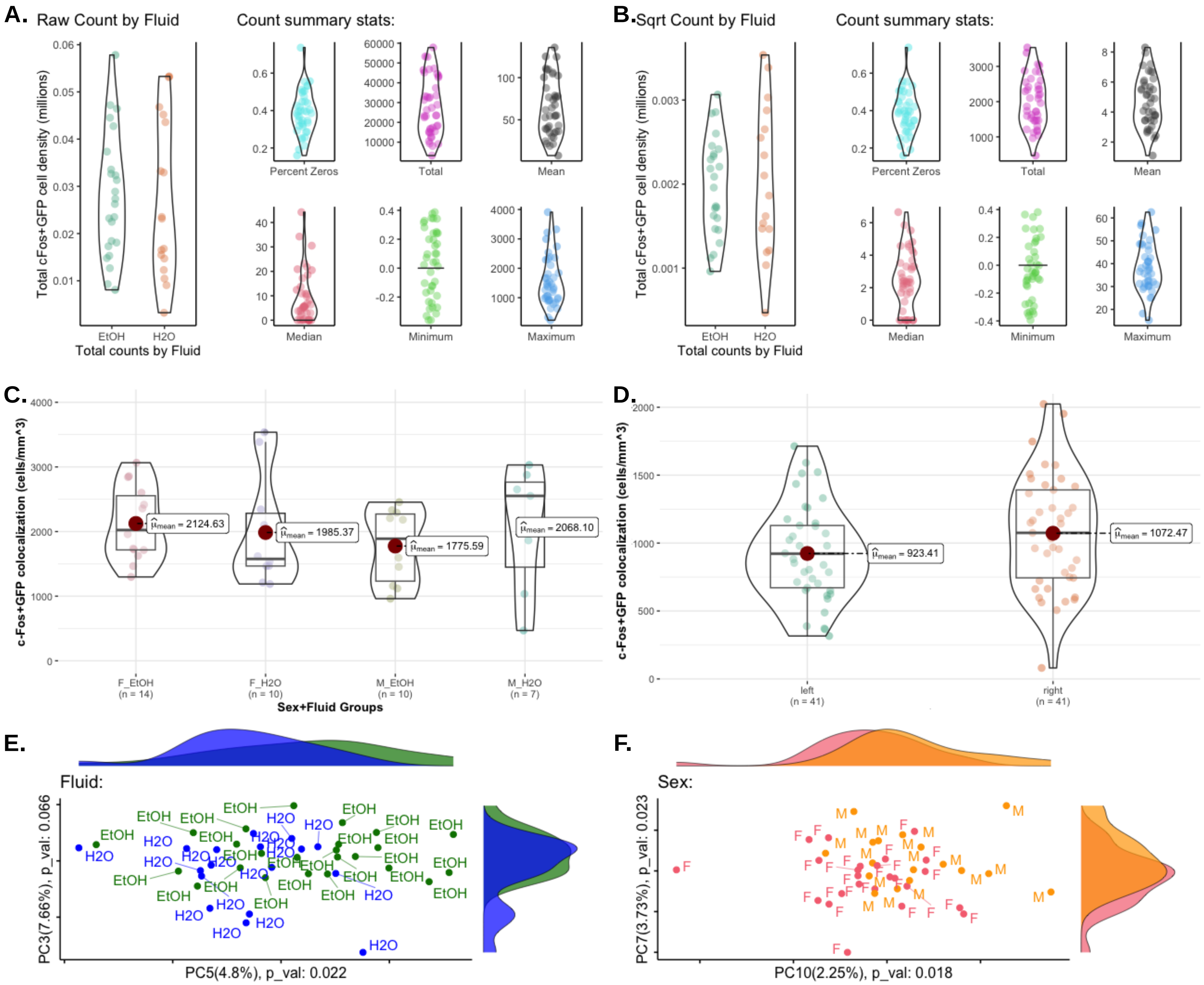
A.
Raw Count by Fluid
Count summary stats:
Total cFos+GFP cell density (millions)
EtOH
H2O
Total counts by Fluid
Percent Zeros
Total
Mean
Median
Minimum
Maximum
B.
Sqrt Count by Fluid
Count summary stats:
Total cFos+GFP cell density (millions)
EtOH
H2O
Total counts by Fluid
Percent Zeros
Total
Mean
Median
Minimum
Maximum
C.
c-Fos+GFP colocalization (cells/mm^3)
$\hat{\mu}_{mean}$ = 2124.63
$\hat{\mu}_{mean}$ = 1985.37
$\hat{\mu}_{mean}$ = 1775.59
$\hat{\mu}_{mean}$ = 2068.10
F_EtOH (n = 14)
F_H2O (n = 10)
M_EtOH (n = 10)
M_H2O (n = 7)
Sex+Fluid Groups
D.
c-Fos+GFP colocalization (cells/mm^3)
$\hat{\mu}_{mean}$ = 923.41
$\hat{\mu}_{mean}$ = 1072.47
left (n = 41)
right (n = 41)
E.
Fluid:
PC3(7.66%), p_val: 0.066
PC5(4.8%), p_val: 0.022
F.
Sex:
PC7(3.73%), p_val: 0.023
PC10(2.25%), p_val: 0.018

Figure S6

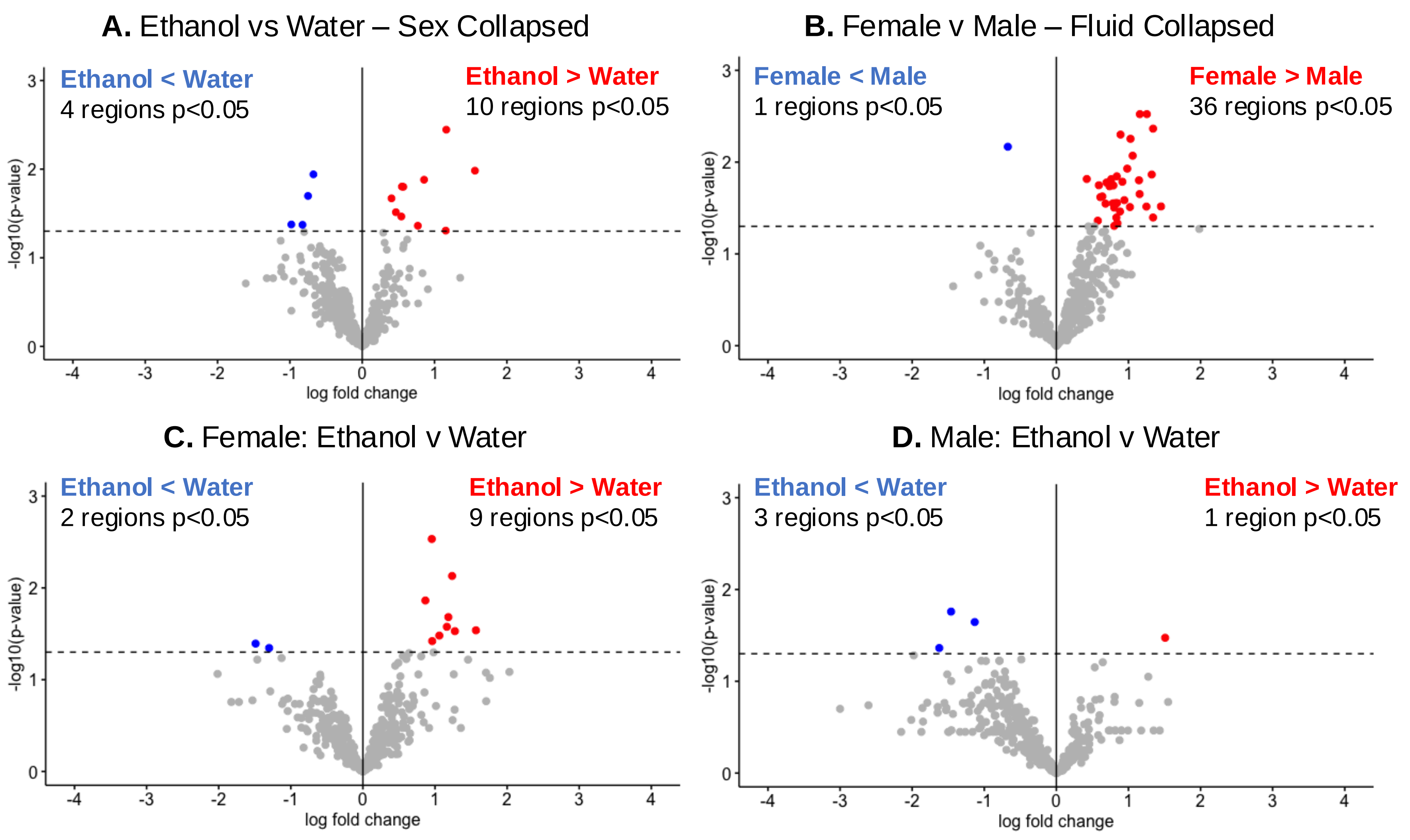
A. Ethanol vs Water – Sex Collapsed
Ethanol < Water
4 regions p<0.05
Ethanol > Water
10 regions p<0.05
B. Female v Male – Fluid Collapsed
Female < Male
1 regions p<0.05
Female > Male
36 regions p<0.05
C. Female: Ethanol v Water
Ethanol < Water
2 regions p<0.05
Ethanol > Water
9 regions p<0.05
D. Male: Ethanol v Water
Ethanol < Water
3 regions p<0.05
Ethanol > Water
1 region p<0.05
-log10(p-value)
log fold change

Figure S7

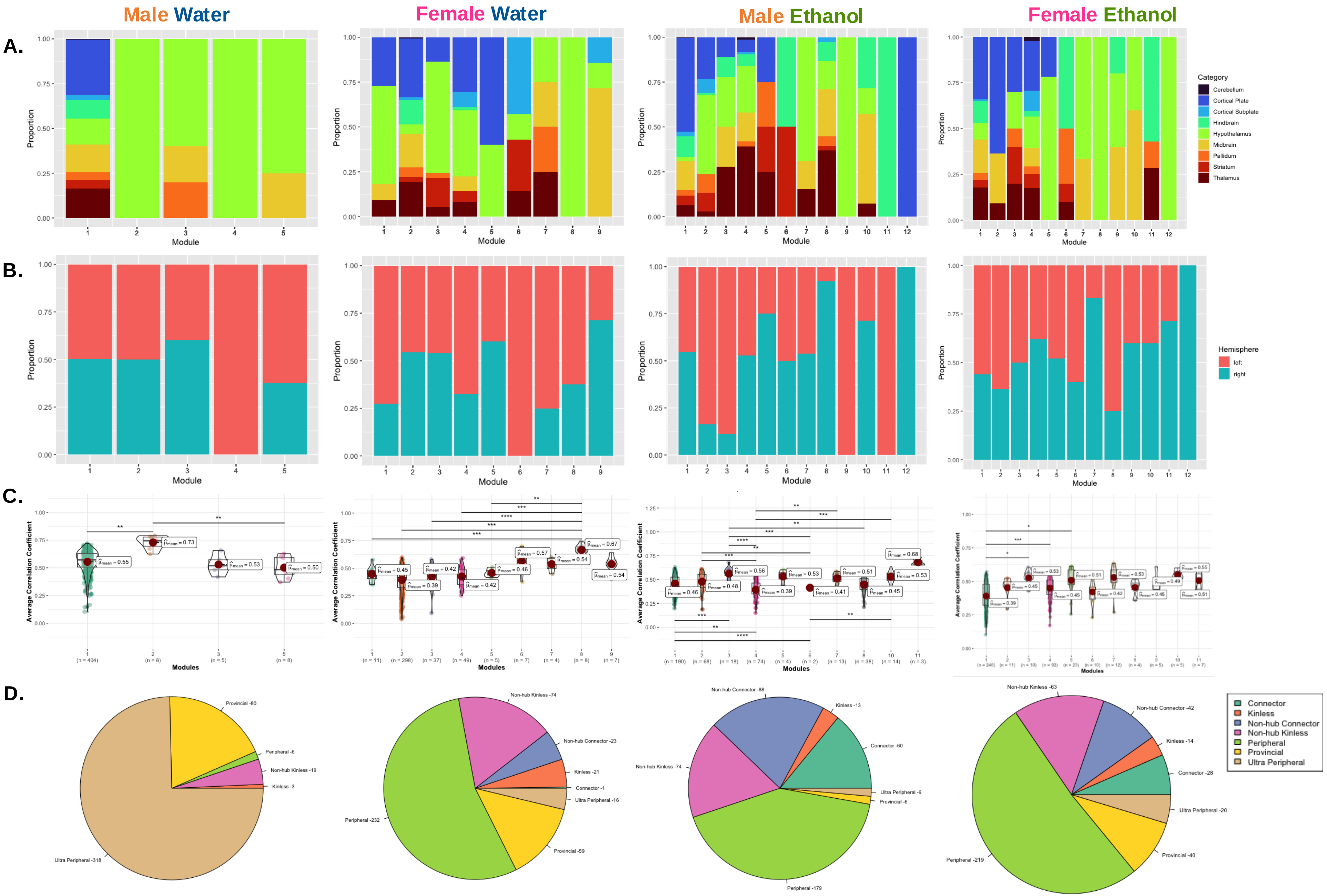

Male Water
Female Water
Male Ethanol
Female Ethanol
A.
B.
C.
D.
Proportion
Module
Category
Cerebellum
Cortical Plate
Cortical Subplate
Hindbrain
Hypothalamus
Midbrain
Pallidum
Striatum
Thalamus
Hemisphere
left
right
Average Correlation Coefficient
Modules
Provincial -80
Peripheral -6
Non-hub Kinless -19
Kinless -3
Ultra Peripheral -318
Non-hub Kinless -74
Non-hub Connector -23
Kinless -21
Connector -1
Ultra Peripheral -16
Provincial -59
Peripheral -232
Non-hub Connector -88
Kinless -13
Connector -60
Ultra Peripheral -6
Provincial -6
Peripheral -179
Non-hub Kinless -74
Non-hub Kinless -63
Non-hub Connector -42
Kinless -14
Connector -28
Ultra Peripheral -20
Provincial -40
Peripheral -219
Connector
Kinless
Non-hub Connector
Non-hub Kinless
Peripheral
Provincial
Ultra Peripheral


Figure S8

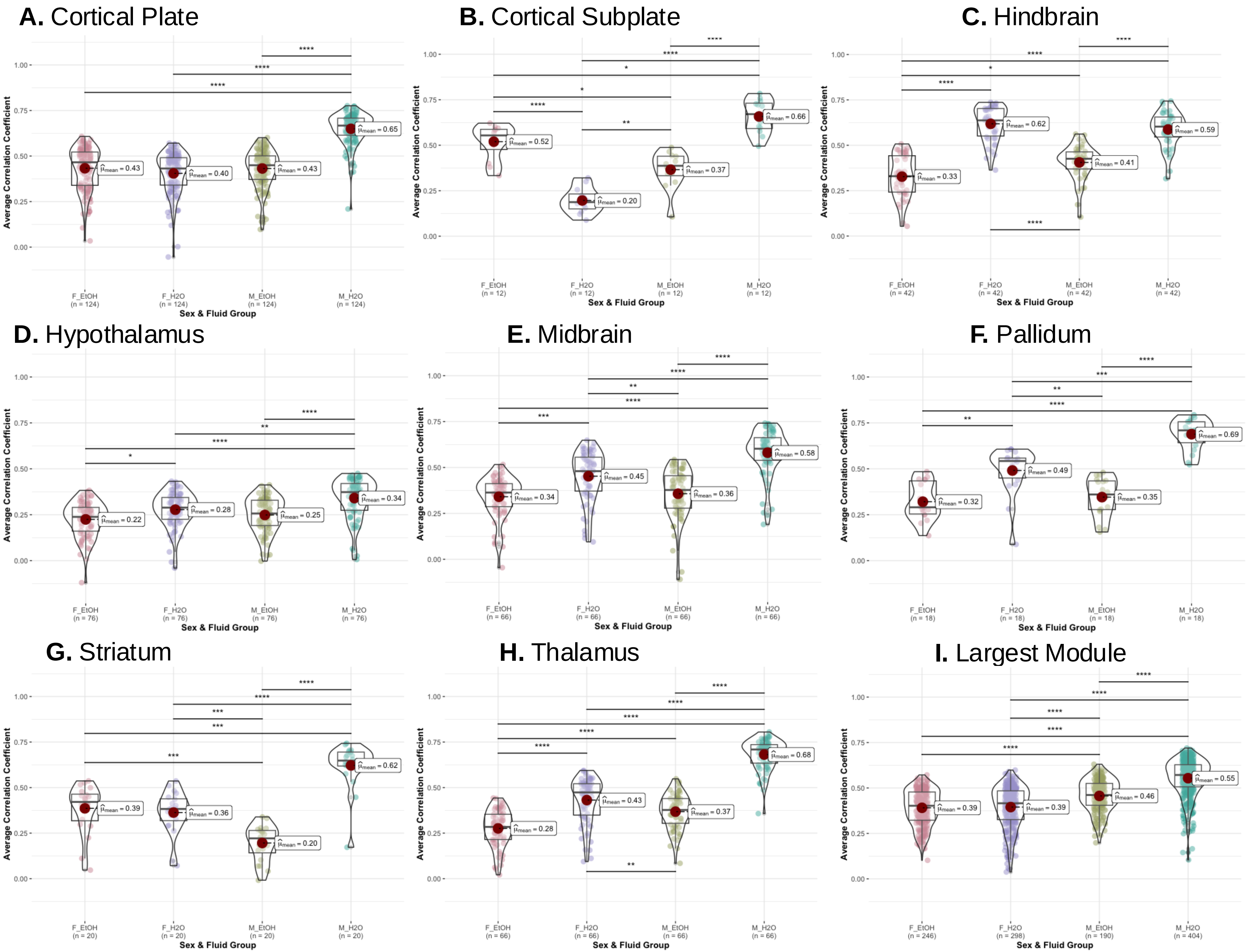
A. Cortical Plate
B. Cortical Subplate
C. Hindbrain
D. Hypothalamus
E. Midbrain
F. Pallidum
G. Striatum
H. Thalamus
I. Largest Module
Average Correlation Coefficient
Sex & Fluid Group
F_EtOH
F_H2O
M_EtOH
M_H2O

Figure S9

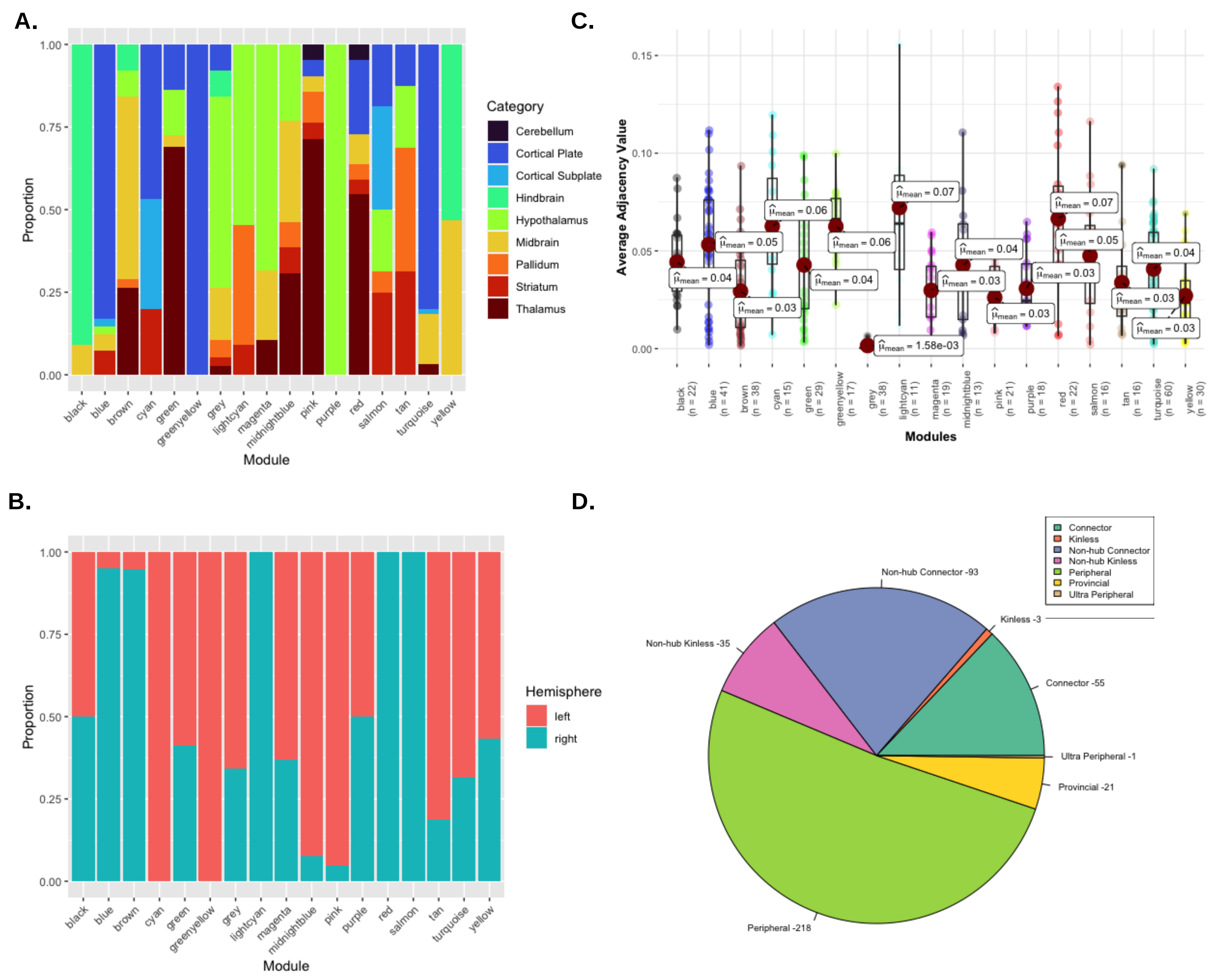


| Significant Module Pair-wise Comparisons | | | |
|---|---|---|---|
| **Group 1** | **Group 2** | **Holm's adj p-val** | |
| black | grey | 1.70e-05 | *** |
| blue | grey | 2.11e-09 | **** |
| brown | greenyellow | 0.02 | * |
| brown | grey | 6.09e-05 | *** |
| cyan | grey | 0.03 | * |
| green | grey | 2.77e-04 | *** |
| greenyellow | grey | 4.19e-06 | **** |
| greenyellow | pink | 6.08e-03 | ** |
| greenyellow | yellow | 5.75e-03 | ** |
| grey | magenta | 0.03 | * |
| grey | pink | 0.01 | * |
| grey | purple | 0.03 | * |
| grey | red | 8.04e-04 | *** |
| grey | turquoise | 2.72e-09 | **** |
| grey | yellow | 3.75e-04 | *** |

Figure S10
